# Harnessing the Power of Antibodies to Fight Bone Metastasis

**DOI:** 10.1101/2021.02.19.432037

**Authors:** Zeru Tian, Ling Wu, Chenfei Yu, Yuda Chen, Zhan Xu, Igor Bado, Axel Loredo, Lushun Wang, Hai Wang, Kuan-lin Wu, Weijie Zhang, Xiang H. -F. Zhang, Han Xiao

**Affiliations:** Department of Chemistry, Rice University, 6100 Main Street, Houston, Texas, 77005; Department of Molecular and Cellular Biology, Baylor College of Medicine, 1 Baylor Plaza, Houston, Texas, 77030; Department of Biosciences, Rice University, 6100 Main Street, Houston, Texas, 77005; Department of Bioengineering, Rice University, 6100 Main Street, Houston, Texas, 77005

## Abstract

Over the past 20 years, antibody-based therapies have proved to be of great value in cancer treatment. Despite the clinical success of these biopharmaceuticals, reaching targets in the bone micro-environment has proved to be difficult perhaps due to the relatively low vascularization of bone tissue and the presence of physical barriers that impair drug penetration. Here, we have used an innovative bone targeting (BonTarg) technology to generate a first-in-class bone-targeting antibody. Moreover, we have used two xenograft models to demonstrate the enhanced therapeutic efficacy of this bone-targeting antibody against bone metastases, compared to the efficacy of traditional antibodies. Our strategy involves the use of pClick antibody conjugation technology to chemically couple the bone-targeting moiety bisphosphonate to the human epidermal growth factor receptor 2 (HER2)-specific antibody trastuzumab. Bisphosphonate modification of therapeutic antibodies results in delivery of higher conjugate concentrations to the bone metastatic niche, relative to other tissues. In both HER2-positive and negative xenograft mice models, this strategy provides enhanced inhibition of experimental bone metastases as well as multi-organ secondary metastases that arise from the bone lesions. Specific delivery of therapeutic antibodies to the bone therefore represents a promising strategy for the treatment of bone metastatic cancers and other bone diseases.

## Introduction

Antibody-based therapies, including those using monoclonal antibodies, antibody-drug conjugates, bispecific antibodies, checkpoint inhibitors, and others, have realized their clinical potential in terms of their power to treat a variety of cancers.^1–4^ Nevertheless, despite the fact that most therapeutic antibodies have high affinities for their targets, the presence of these same targets in normal tissues can dramatically limit the ability of therapeutic agents to hit their targets without inducing unacceptable “on-target” toxicity in healthy cells.^5–7^ Furthermore, low levels of delivery of therapeutic antibodies to some tissues such as brain or bone can significantly limit their efficacy in treating diseases in these tissues.^8^ Thus, it is likely that enhancing both the antigen and tissue specificity of antibodies will ultimately transform the efficacy of antibody therapy for clinical treatment of cancer.

Half of patients with an initial diagnosis of metastatic breast cancer (BCa) will develop bone metastases.^9^ Patients having only skeletal metastases usually have a better prognosis than patients with vital organ metastases.^9,10^ Furthermore, bone metastasis is associated with severe symptoms such as spinal cord compression, pathological fractures, and hypercalcemia.^11^ Despite our deep understanding of molecular mechanisms,^12,13^ effective therapies that can eliminate cancer cells are still lacking.^14^ Bone is not the final destination of metastatic dissemination. Recent genomic analyses have revealed frequent “metastasis-to-metastasis” seeding.^15–17^ Over two-thirds of bone-only metastases subsequently develop secondary metastases to other organs, ultimately leading to the death of patients.^9,10^ In fact, some metastases initially identified in non-bone organs are actually the result of seeding from sub-clinical bone micrometastases (BMMs). This apparently is the result of cancer cells initially arriving in the bone and then acquiring more aggressive phenotypes that allow them to establish more overt metastases in both bone and other sites.^18^ It should therefore be useful to develop strategies for preventing BMMs from establishing more overt metastases in both bone and non-bone tissues.

While targeted antibody therapy and immunotherapy are currently emerging as new avenues for treating metastatic breast cancer, the performance of these agents in patients with bone metastases has been disappointing. For example, trastuzumab (Herceptin) and pertuzumab (Perjeta) antibodies targeting human epidermal growth factor receptor 2 (HER2) have been used to treat patients in adjuvant and metastatic settings. Although many BCa patients benefit from these treatments, in large numbers of BCa patients with bone metastasis, the disease progresses within one year and few patients experience prolonged remission.^19–22^ In another phase III clinical trial testing atezolizumab in patients with metastatic triple-negative BCa, progression-free survival was significantly longer in the atezolizumab group than in the placebo group. However, among BCa patients with bone metastases, no significant difference was observed between the atezolizumab-treated and placebo groups for risk of progression or death.^23^ Therapies with improved outcomes for BCa patients with bone metastases are therefore highly desired.

Attempts to ensure effective concentrations of a therapeutic drug in bone unavoidably lead to high concentrations in other tissues as well, often resulting in adverse systemic effects or side effects that may limit or exclude the use of the drug.^24,25^ In this case, the potential benefit of passive targeting is lost. Here, we describe an innovative bone targeting (BonTarg) technology that enables the tissue-specific delivery of therapeutic antibodies to the bone via conjugation of bone-targeting moieties. The resulting bone-targeting antibodies can specifically target the bone metastatic niche to eliminate bone micrometastases and also prevent seeding of multi-organ metastases from bone lesions. Taking advantage of the high mineral concentration unique to the bone hydroxyapatite matrix, bisphosphonate (BP) conjugation has been used for selective delivery of small molecule drugs, imaging probes, nuclear medicines, and nanoparticles to the bone as a means of treating of osteoporosis, primary and metastatic bone neoplasms, and other bone disorders.^24,26–30^ Negatively-charged BP has a high affinity for hydroxyapatite (HA), which is the main component of hard bone, resulting in preferential binding to the bone. However, the potential benefit of bone-specific delivery of large therapeutic proteins to the bone by modifying BP hasn’t yet been explored. We have used pClick conjugation technology to site-specifically couple the BP drug Alendronate (ALN) to the HER2-targeting monoclonal antibody trastuzumab (Tras).^31^ In two xenograft models based on intra-iliac artery (IIA) injection, the resulting trastuzumab-Alendronate conjugate (Tras-ALN) significantly enhances the concentration of therapeutic antibody in the bone metastatic niche, inhibits cancer development in the bone, and limits secondary metastases to other organs. This type of specific delivery of therapeutic antibodies to the bone has the potential to enhance both the breadth and potency of antibody therapy for bone-related diseases.

## Results

### Development of the First Bone-Targeting Antibody using BonTarg

To explore the possibility of specifically delivering therapeutic antibodies to the bone via conjugation to BP molecules, we designed a model using the HER2 targeting antibody trastuzumab (Tras) and the BP drug Alendronate (ALN). ALN is a second-generation BP drug that is used as a bone-targeting agent as well as a regimen for treating osteoporosis and bone metastasis.^31^ To ensure that ALN conjugation does not impair the therapeutic efficacy of the antibody, we have employed a novel proximity-induced antibody conjugation strategy named pClick.^31^ pClick technology enables the site-specific attachment of payloads to native antibodies under mild conditions, thus minimizing the disruption of binding to the antigen receptor or the FcγRIII receptor, the receptor responsible for activating antibody-dependent cell-mediated cytotoxicity (ADCC). The pClick technology does not rely on antibody engineering or on the UV/chemical/enzymatic treatments that characterize the generation of most therapeutic antibodies. To prepare trastuzumab-Alendronate conjugates (Tras-ALN), we first used pClick to generate Tras containing an azide functional moiety, followed by reaction with bicyclo[6.1.0]nonyne (BCN)-functionalized ALN (**Fig. 1A, and S1-S3**). The resulting Tras-ALN was further purified on a desalting column and fully characterized by SDS-PAGE and ESI-MS (**Fig. 1B, C**). To our delight, no unconjugated heavy chain or degradation products were revealed by SDS-PAGE, indicating a more than 95% coupling efficiency. ESI-MS analysis also revealed that more than 95% of the heavy chain was conjugated with the ALN molecule.

**Fig 1.**
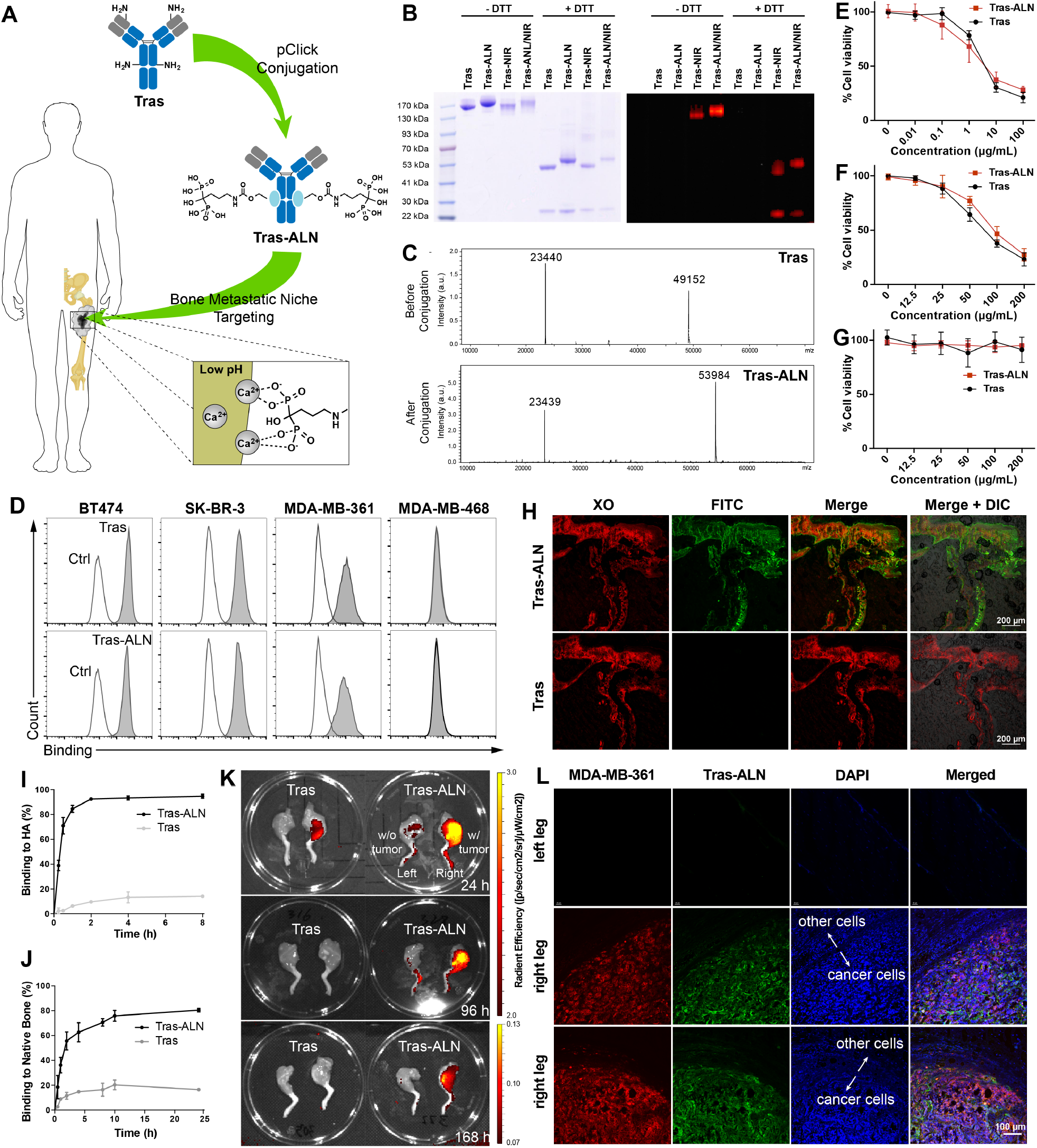
**(A)** Therapeutic antibodies can be site-specifically delivered to bone by pClick conjugation of bisphosphonate molecules that bind to the bone hydroxyapatite matrix. **(B)** SDS-PAGE analysis of Tras, Tras-ALN, and their near-infrared (NIR) fluorophore conjugates under reducing and non-reducing conditions, visualized by coomassie blue staining (left) and a fluorescence scanner (right) **(C)** Mass spectrometry analysis of Tras and Tras-ALN. **(D)** Flow cytometric profiles of Tras and Tras-ALN binding to BT474 (HER2+++), SK-BR-3 (HER2+++), MDA-MB-361 (HER2++), and MDA-MB-468 (HER2-) cells. **(E-G)** *In vitro* cytotoxicity of Tras and Tras-ALN against BT474, MDA-MB-361, and MDA-MB-468 cells. **(H)** Differential bone targeting ability of unmodified Tras and Tras-ALN conjugate. Nondecalcified bone sections from C57/BL6 mice were incubated with 50 µg/mL Tras or Tras-ALN overnight, followed by staining with fluorescein isothiocyanate (FITC)-labeled anti-human IgG and 4 µg/mL xylenol orange (XO, known to label bone), Scale bars, 200 µm. **(I-J)** Binding kinetics of Tras and Tras-ALN to hydroxyapatite (HA) and native bone. **(K)** *Ex vivo* fluorescence images of lower limbs of athymic nude mice bearing MDA-MB-361 tumors 24 h, 96 h, or 168 h after the retro-orbital injection of Cy7.5-labeled Tras and Tras-ALN. Tumor cells were inocu6lated into the right limbs of nude mice via IIA injection. **(L)** Nondecalcified bone sections from the biodistribution study were stained with FITC-labeled anti-human IgG (green), RFP (red) and DAPI (blue), Scale bars, 100 µm.

### Antibody Conjugation to ALN Retains Antigen Binding and Specificity

To investigate the effect of ALN conjugation on antigen-binding affinity and specificity, binding affinities of Tras and Tras-ALN were assessed by flow cytometry analysis of HER2-positive and negative cell lines. Fig. 1D reveals that both Tras and Tras-ALN have strong binding affinities for the HER2-expressing cell lines BT474, SK-BR-3, and MDA-MB-361, but not for the HER2-negative cell line MDA-MB-468, suggesting that the antibody specificity was not altered by ALN conjugation (**Table S1**). The Kd values for binding to HER2-positive cells are within a similar range for Tras and Tras-ALN (BT474, 3.0 vs 3.8 nM; SK-BR-3, 2.3 vs 3.0 nM, respectively), indicating that ALN conjugation does not affect the strength of antigen-binding (**Fig. S4-S7**). Confocal fluorescent imaging further confirms that Tras-ALN retains antigen binding and specificity (**Fig. S8**). HER2-positive BT474 and SK-BR-3 cells, and HER2-negative

MDA-MB-468 cells were incubated for 30 mins with fluorescein isothiocyanate (FITC)-labeled Tras-ALN. Confocal imaging indicates that cell-surface-associated fluorescence is only exhibited for HER2-positive BT474 and SK-BR-3 cells, and not for HER2-negative MDA-MB-468 cells (**Fig. S8**). Thus, ALN modification of Tras does not affect its antigen-binding affinity and specificity. Next, the Tras-ALN conjugate was tested for selective cytotoxicity against HER2-expressing and HER2-negative breast cancer cells. As shown in **Fig. 1E, 1F, 1G** and **Table S1**, the Tras-ALN conjugate exhibits cytotoxic activity against HER2-positive BT-474 cells (EC50 of 2.3 ± 0.7 µg/ml) and MDA-MB-361 (EC50 of 78 ± 21 µg/ml) that is indistinguishable from that of Tras (EC50 of 1.4 ± 0.9 µg/ml and EC50 of 57 ± 10 µg/ml). Neither antibody kills HER2-negative MDA-MB-468 cells (EC50 >500 µg/ml). These results indicate that conjugation of the negatively charged moiety ALN preserves the antigen-binding and *in vitro* anti-tumor cell activity of the Tras antibody.

### Enhanced Targeting of the Bone Metastatic Niche by Tras-ALN *in vitro* and *in vivo*

We next explored the ability of the Tras-ALN conjugate to target bone tissue. Non-decalcified bone sections from C57BL/6 mice were incubated overnight at 4°C with 50 µg/mL Tras or Tras-ALN conjugate, followed by labeling with FITC-labeled anti-human IgG. Before imaging via confocal laser scanning microscopy, these bone sections were further stained for 30 min with 4 µg/mL xylenol orange (XO, known to label bone). We observed a FITC signal in sections stained with the Tras-ALN conjugate, but not in sections stained with unmodified Tras (**Fig. 1H**). Furthermore, localization of the Tras-ALN signal correlated well with the XO signal, confirming the specific targeting of bone by Tras-ALN. To quantify the difference in affinity between binding of the Tras-ALN conjugate and unmodified Tras, we incubated Tras-ALN and Tras with hydroxyapatite or native bone. As shown in **Fig. 1I and 1J**, unmodified Tras exhibited only slight binding to HA or native bone. Even with an increase in the incubation time, the binding affinity of Tras did not change significantly. In contrast, approximately 80%-90% of Tras-ALN was bound to HA and native bone after 2 h and 10 h, respectively.

Encouraged by the *in vitro* bone-targeting ability of ALN-conjugated Tras, we carried out an *in vivo* biodistribution study with the Tras-ALN conjugate using a tumor xenograft model. To facilitate the detection of antibodies *in vivo*, we first conjugated Tras and Tras-ALN with Cyanine 7.5 (Cy7.5)-hydroxysuccinimide (NHS) ester. The resulting Cy7.5 labeled conjugates were analyzed using SDS-PAGE. As expected, fluorescence was associated only with the Cy7.5-labeled conjugates (**Fig. 1B**). An important feature of BP is that uptake of bisphosphonate into bone metastases is much higher than in healthy bone tissue, due to the relatively low pH of the bone metastatic microenvironment.^32–35^ To investigate if ALN-Tras can specifically target bone metastases, thus minimizing on-target toxicity to normal bone tissue, we evaluated the targeting properties of ALN-Tras in a bone tumor model. We created a bone micrometastasis model by using intra-iliac artery (IIA) injection of MDA-MB-361 cells labeled with luciferase and red fluorescent protein (RFP) into the right hind limbs of nude mice. IIA injection is a novel technology recently developed in our lab for establishing bone micrometastases. Our method allows for selective delivery of cancer cells into hind limb bones without causing tissue damage.^36–38^ This technology allows sufficient time for some indolent cells to eventually colonize the bone as well as a large number of cancer cells to specifically colonize the bone, thereby enriching micrometastases in early stages. This allows for swift detection and robust quantification of micrometastases. Establishment of micrometastases was followed by treatment with Tras or Tras-ALN (1 mg/kg). 24, 96 or 168 hrs after administration of antibody or antibody conjugate, the major organs, including heart, liver, spleen, kidney, lung, and bone, were removed and analyzed using the Caliper IVIS Lumina II imager (**Fig. 1K and S9**). Significantly, *ex vivo* fluorescence images at 96 h post-injection of antibody confirmed clear accumulation of Cy7.5-labeled Tras-ALN in the bone compared with Cy7.5-labeled Tras (**Fig. 1K and S10**). Furthermore, the uptake of Tras-ALN into cancer-bearing bones is significantly higher than into healthy bone tissue. This is consistent with previous observations that BP molecules prefer to bind to the bone matrix in an acidic tumor environment.^39^ In a separate study, unlabeled Tras-ALN (1 mg/kg) was administered into the nude mice bearing MDA-MB-361 tumor in the right hind limb. Bone sections from this study were also stained with FITC-labeled anti-human IgG, RFP and DAPI. We only observed FITC signals in sections from the right leg harboring MDA-MB-361 tumors. No FITC signals were detected in the left leg without tumors (**Fig. 1L**). Significantly, the FITC signal correlated well with the red fluorescence of MDA-MB-361 cells, suggesting that Tras-ALN conjugate selectively targets the bone metastatic site, but not healthy bone. These results demonstrate that ALN conjugation can significantly enhance the delivery and concentration of therapeutic antibodies in bone metastatic sites.

### Enhanced Therapeutic Efficacy of Tras-ALN Against Bone Micrometastases

To determine whether bone-targeting trastuzumab represents a novel therapeutic approach for treating micrometastases of breast cancer in the bone, we carried out a xenograft study in nude mice. Using intra-iliac artery (IIA) injection, we inoculated the right hind limbs of nude mice with 5 x 10^5^ MDA-MB-361 cells labeled with firefly luciferase. Five days after the IIA injections, mice were treated with phosphate-buffered saline (PBS), ALN (10 µg/kg), Tras (1 mg/kg), or Tras-ALN (1 mg/kg) via retro-orbital injection. As shown in **Fig. 2A and S11**, micrometastases in PBS- and ALN-treated mice accumulated rapidly, while development of lesions in Tras- and Tras-ALN-treated mice was delayed. Whole-body bioluminescence imaging (BLI) signals suggested that treatment with Tras-ALN resulted in more significant inhibition of micrometastasis progression, compared to that seen in Tras-treated mice (**Fig. S12A and S12B**). The increases in BLI from day 6 to 87 showed that the Tras-ALN-treated group had fewer fold-increases in the tumor sizes compared to Tras-treated group (Tras vs Tras-ALN: 1965.1 ± 798.3 vs 42.6 ± 23.4, **Fig. 2B and 2C**). As we built the bone metastasis in the hind limbs, the effect of Tras-ALN on the BLI signal in the hind limbs was also quantified. Similar to whole-body BLI signal, Tras-ALN-treated group had less BLI signal intensity and fewer fold-increase in the hind limbs **(Fig. S13)**. Moreover, survival of Tras-ALN-treated mice was notably enhanced compared to that of PBS-, ALN-, and Tras-treated mice, demonstrating the efficacy of Tras-ALN against HER2-positive cells *in vivo* (**Fig. 2D**). Furthermore, no weight loss as a sign of ill health was observed in any of the treated mice, suggesting the absence of toxicity associated with the bone-targeting antibodies (**Fig. 2E)**.

**Fig 2.**
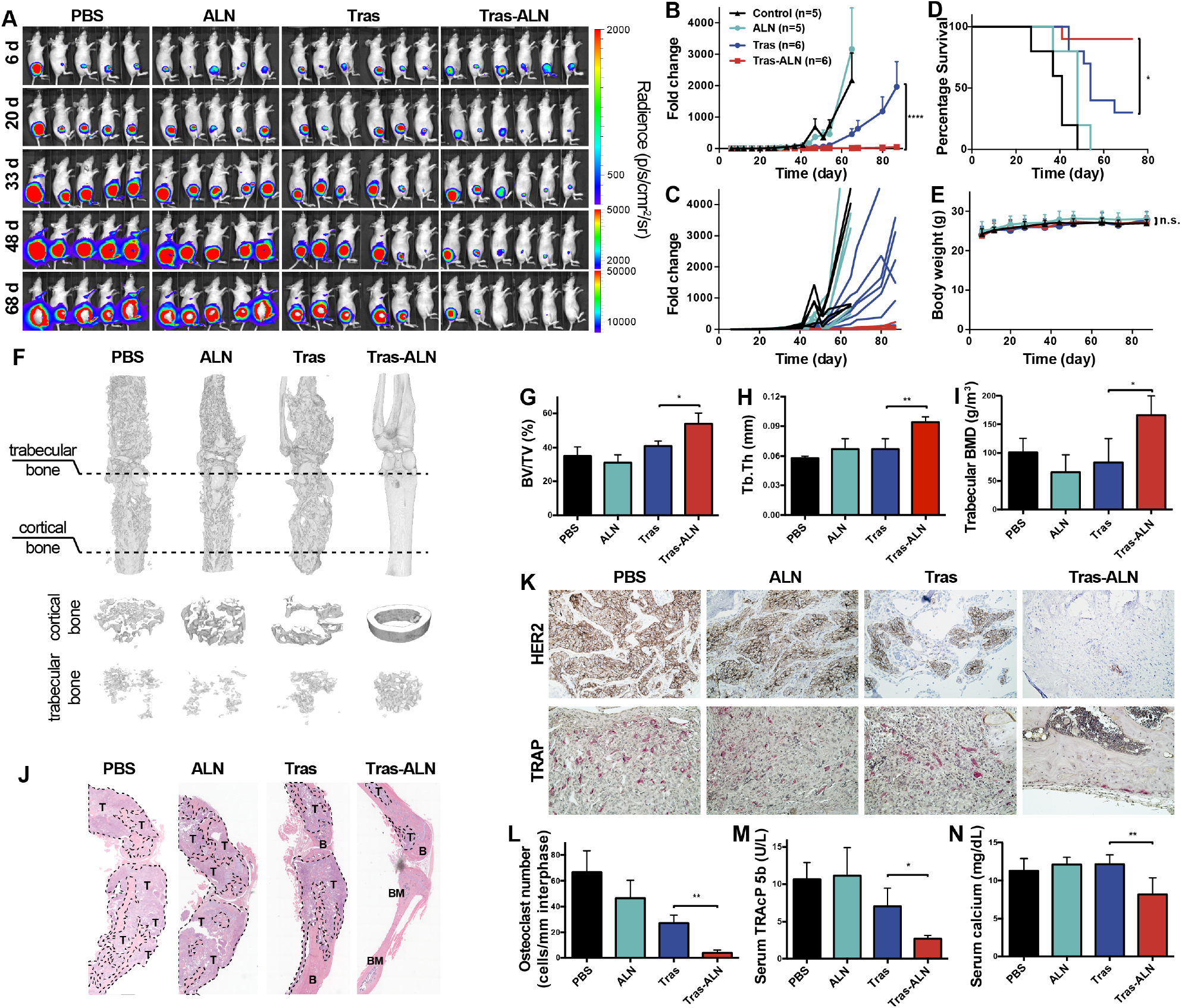
Tras-ALN inhibits breast cancer micrometastases in the bone. **(A)** MDA-MB-361 cells were IIA injected into the right hind limb of nude mice, followed by treatment with PBS, ALN (10 µg/kg retro-orbital venous sinus in PBS twice a week), Tras (1 mg/kg retro-orbital venous sinus in sterile PBS twice a week), and Tras-ALN conjugate (same as Tras). Tumor burden was monitored by weekly bioluminescence imaging. **(B)** Fold-change in mean luminescent intensity of MDA-MB-361 tumors in mice treated as described in (A), two-way ANOVA comparing Tras to Tras-ALN. **(C)** Fold-change in Individual luminescent intensity of HER2-positive MDA-MB-361 tumors in mice treated as described in (A). **(D)** Kaplan-Meier plot of the time-to-euthanasia of mice treated as described in (A). For each individual mouse, the BLI signal in the whole body reached 10^7^ photons sec^−1^ was considered as the endpoint. **(E)** Body weight change of tumor-bearing mice over time. **(F)** MicroCT scanning in the supine position for groups treated with PBS, ALN, Tras, or Tras-ALN 82 days after tumor implantation. **(G)** Quantitative analysis of bone volume density (BV/TV). **(H)** Quantitative analysis of trabecular thickness (Tb.Th). **(I)** Quantitative analysis of trabecular bone mineral density (BMD). **(J)** Representative longitudinal, midsagittal hematoxylin and eosin (H&E)-stained sections of tibia/femur from each group. T: tumor; B: bone; BM: bone marrow. **(K)** Representative images of HER2 and TRAP staining of bone sections from each group. **(L)** Osteoclast number per image calculated at the tumor-bone interface in each group (pink cells in (K) were considered as osteoclast positive cells). Serum TRAcP 5b levels of mice treated as described in (A). **(N)** Serum calcium levels of mice treated as described in (A). _****_*P* < 0.0001, _***_*P* < 0.001, _**_*P* < 0.01, _*_*P* < 0.05, and n.s. = *P* > 0.05.

These results were further confirmed by micro-computed tomography (microCT) data and histology, emphasizing the finding that bone-targeting antibodies can decrease both the number and the extent of osteolytic lesions. As shown in **Fig. 2F and Fig. S14**, femurs from PBS-, ALN-, and Tras-treated groups exhibited significant losses of bone mass, while bone loss in the Tras-ALN-treated group was much reduced. Quantitative analysis revealed that the Tras-ALN-treated group had significantly higher bone volume (**Fig. 2G**,. 6B: BV/TV (%), 35.08 ± 2.65 vs 56.67 ± 1.02, p=0.0005) thicker trabecular bone (**Fig. 2H**, Tb.Th (mm), 0.061 ± 0.003 vs 0.094 ± 0.002, p=0.003), and higher trabecular bone mineral density (**Fig. 2I**, BMD (mg/mm^3^), 101.16 ± 12.24 vs 165.94 ± 12.84, p=0.035) compared to the Tras-treated group.

Tumor size was also analyzed by histomorphometric analysis of the bone sections. Tibiae and femurs from the PBS-treated and ALN-treated groups had high tumor burdens (**Fig. 2J**). Tras treatment slightly reduced the tumor burden, but the reduction was not statistically significant. In contrast, a significant reduction of tumor burden was observed in the Tras-ALN-treated group. Histological examination of the bone samples from various treatment groups reveals that bone matrix is generally destroyed in bones with high tumor burden, whereas bones with less tumor burden in the Tras-ALN-treated group exhibit intact bone matrix. The reduction of tumor burden was also confirmed by HER2 immunohistochemistry (IHC). As shown in Fig. 2K, the number of HER2-positive breast cancer cells is dramatically decreased in Tras-ALN-treated mice, even though HER2 expression by individual tumor cells is unchanged. This suggests that extended treatment with Tras-ALN has no effect on HER2 expression by MDA-MB-361 cells.

To examine Tras-ALN inhibition of tumor-induced osteolytic bone destruction, we examined the bone-resorbing, tartrate-resistant, acid phosphatase-positive multinucleated osteoclasts in bone samples (**Fig. 2K**). Tartrate-resistant acid phosphatase (TRAP) staining identified reduced numbers of osteoclasts (pink cells) lining the eroded bone surface in Tras-ALN-treated mice, compared to Tras-treated mice (**Fig. 2K, L, and S15**). Serum TRAcP 5b and calcium levels, indicators of bone resorption, were also measured at the experimental endpoint. Significantly higher reductions in bone resorption were observed in the Tras-ALN-treated group (**Fig. 2M and N**). Taken together, these results indicate that bisphosphonate modification of therapeutic antibodies significantly enhanced their ability to retard the development of micrometastasis-induced osteolytic lesions (**Table S2**).

### Tras-ALN inhibits multi-organ metastases from bone lesions

In more than two-thirds of cases, bone metastases are not confined to the skeleton, but rather give rise to subsequent metastases to other organs.^9,10^ While we have used IIA injection to investigate early-stage bone colonization, as these bone lesions progress over an 8-12 week period, metastases begin to appear in other organs, including additional bones, lungs, liver, kidney, and brain. Hence, we investigated the ability of Tras-ALN to reduce the metastasis of HER2-positive MDA-MB-361 cancer cells to other organs. As before, 5 x 10^5^ MDA-MB-361 cells labeled with firefly luciferase were introduced into the right hind limbs of nude mice via IIA injection, followed by treatment with Tras (1 mg/kg) and Tras-ALN (1 mg/kg). Then, mice were subjected to whole-body BLI twice a week following tumor-cell injection. The whole-body and hind limbs BLI signals werequantified and showed in **Fig. S16A**. Secondary metastases in various organs were calculated as follows: BLI signal in whole body – BLI signal in hind limbs. As shown in **Fig. S16**, There was a time-dependent increase in the organs BLI signal to 10^6^ photons sec^−1^ in the Tras treated group. And there was significant inhibition of BLI signal accumulation in organs of Tras-ALN-treated group (P<0.0001). At the endpoint of the study, mice were euthanized, and organs were harvested for bioluminescence imaging. Much higher levels of right hind limb (100%), heart (20%), liver (80%), spleen (40%), lung (60%), kidney (60%) and brain metastasis (40%) were observed in the Tras treated group, compared to the right hind limb (42.9%) and liver (14.3%, **Fig. 3A, B**, and **S17**) in the Tras-ALN group. Other organs such as the lungs, spleen, kidney, and brain were devoid of metastases in Tras-ALN-treated mice. Our data indicated that bone-targeting antibodies, compared to unmodified antibodies, can significantly inhibit multi-organ metastases resulting from the dissemination of initial bone micrometastases. Mice treated with Tras-ALN exhibited fewer metastases to other organs than mice in the other treatment groups, establishing the ability of bone-targeting antibodies to inhibit “metastasis-to-metastasis seeding”.

**Fig 3.**
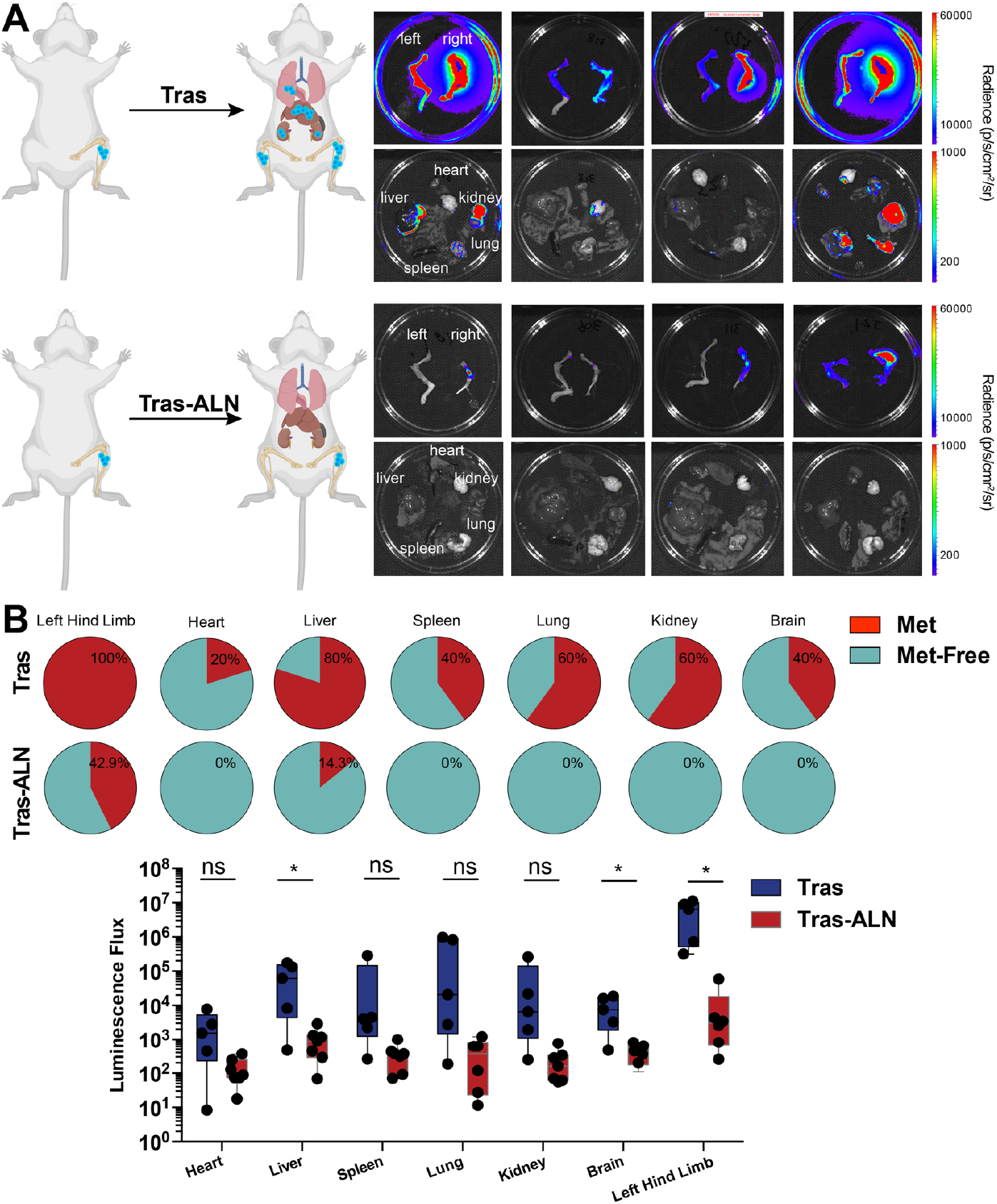
**(A)** Secondary metastases observed in various organs in mice treated with Tras (top) or Tras-ALN (bottom). **(B)** Pie charts show the frequencies of metastasis observed in various organs in mice treated with Tras (1 mg/kg retro-orbitally in sterile PBS twice a week), and Tras-ALN conjugate (same as Tras). **(C)** Quantification of bioluminescence signal intensity in different organs, including other bones, as measurement of metastases resulted from Tras and Tras-ALN-treated mice. p values are based on one-way ANOVA test. _*_*P* < 0.05 and n.s. = *P* > 0.05.

### Enhanced Therapeutic Efficacy of Tras-ALN in a HER2-negative model

Previous reports indicate that a substantial portion of the minimal residual disease seen in HER2-negative patients may nevertheless be due to HER2 signaling^40,41^. It was also reported that HER2 signaling may mediate stem cell properties in a subpopulation of HER2-negative cells, this raises the possibility that anti-HER2 treatment may be able to eradicate bone metastases of both HER2-positive and negative breast cancer.^42^ We therefore evaluated the therapeutic effects of Tras-ALN using breast cancer cells that are not HER2-positive but exhibit HER2 up-regulation specifically in bones. We used intra-iliac artery (IIA) injection to deliver MCF-7 (HER2-, ER+) cancer cells into hind limb bones,^36,38^ followed by treatment with Tras or Tras-ALN (7 mice per group, 1 mg/kg). Mice were imaged twice a week and signal intensity of whole-body and hind limbs and were quantified. As shown in **Fig. 4, S18 and S19**, treatment with Tras-ALN resulted in more significant inhibition of tumor growth than seen in Tras-treated mice, demonstrating the efficacy of Tras-ALN against HER2-negative cells *in vivo* (p<0.005). Meanwhile, significant reductions of serum TRACP 5b (4.41 ± 1.12 U/L, p<0.05) and serum calcium (10.36 ± 0.53 mg/dL, p<0.05) levels were observed in Tras-ALN-treated group (**Fig. S20**). Similar to HER2+ model, secondary metastases in various organs were also exhibited significant reductions in BLI signal (P<0.0001) over the course of the study (**Fig. S21**). Next, we also evaluated the ability of Tras-ALN to inhibit multi-organ metastases from bone lesions *ex vivo*. At day 68, metastatic cells were observed in the right hind limb (83.4%), liver (33.4%), lung (83.4%), and brain (66.7%) in the Tras-treated group, compared to values found in the right hind limb (50%), lung (50%) and brain (50%, **Fig. S22**) of Tras-ALN treated mice. These data suggest that the bone-targeting Tras-ALN conjugate may be useful in preventing the progression of HER2-negative bone micrometastases to overt bone metastases, as well as blocking the secondary metastasis of HER2-negative cells to other organs (**Table S3**).

**Fig 4.**
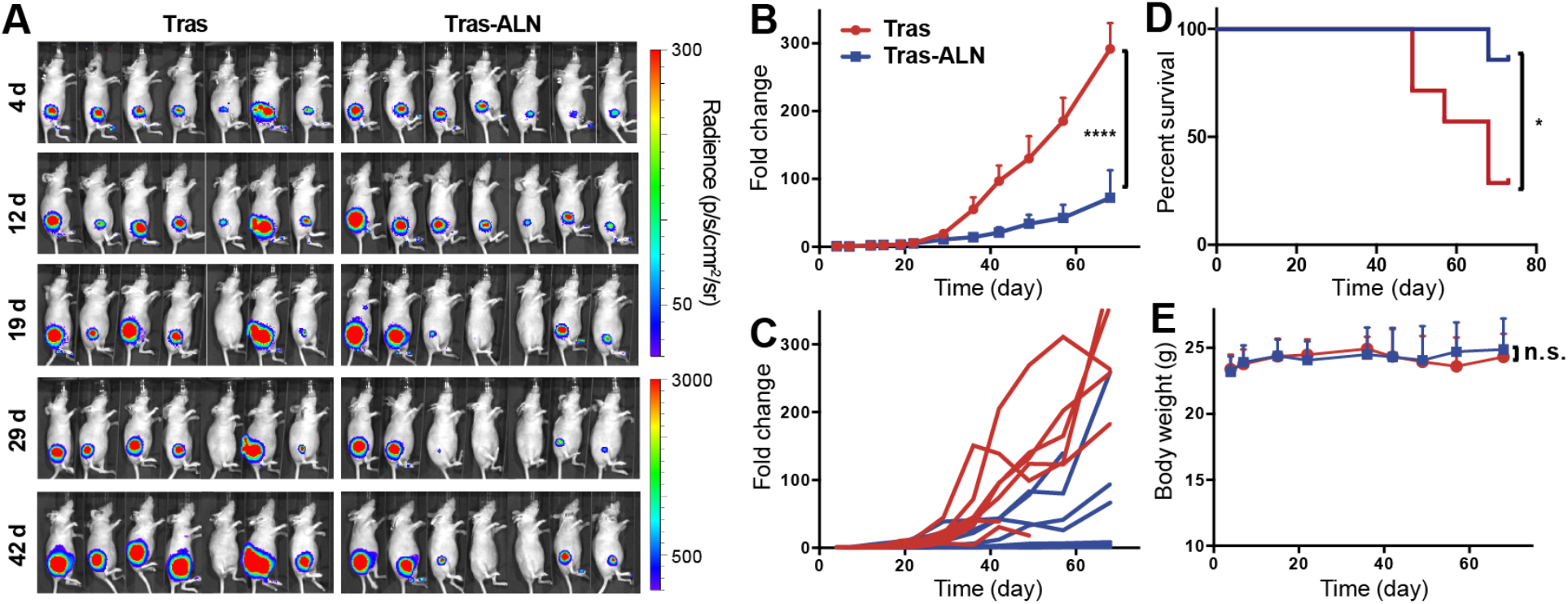
*In vivo* comparison of Tras and Tras-ALN in HER2-negative model. **(A)** Tumor burden was monitored by weekly bioluminescence imaging, and **(B)** quantified by the radiance detected in the region of interest. **(C)** Fold-change in Individual luminescent intensity of HER2-negative MCF-7 tumors in mice treated as described in (A). **(D)** Kaplan-Meier plot of the time-to-sacrifice of mice treated as described in (A). For each individual mouse, the BLI signal in the whole body reached 10^7^ photons sec^−1^ was considered as the endpoint. **(E)** Body weight change of tumor-bearing mice over time. _****_*P* < 0.0001, _*_*P* < 0.05, and n.s. = *P* > 0.05.

## DISCUSSION

Despite the fact that breast cancer (BCa) patients have an extremely good chance of recovery from the disease, 20-40% of BCa survivors will eventually suffer metastases to distant organs.^43^ Metastasis to bone occurs in about 70% of these cases.^44,45^ BCa patients with bone metastases suffer from pain and immobility, along with susceptibility to skeletal-related events (SREs) such as fracture, bone pain, spinal cord compression, and hypercalcemia. SREs significantly reduce quality of life and increase mortality. The one-year survival rate of BCa patients with bone metastases is 51%, but the five-year survival rate drops to 13%.^46,47^ In cases where the skeleton is the only site of metastasis, patients usually have better prognoses than patients with visceral organ metastases.^9,10^ In more than two-thirds of cases, bone metastases will not remain confined to the skeleton, but instead are responsible for subsequent metastases to other organs and eventually to the death of patients.^9,10^ Recent genomic analyses suggest that the majority of metastases are the result of seeding from other metastases, rather than from primary tumors.^15–17^ Some metastases initially found in non-skeletal organs also appear to be seeded from sub-clinical bone micrometastases (BMMs), as suggested by the finding that, subsequent to colonization of bone, metastatic cancer cells in BMMs can acquire more aggressive phenotypes even before establishing overt bone metastases.^18^ Thus, strategies for inhibiting progression of BMMs can prevent further BCa metastasis within the bone, as well as secondary metastases from bone to other organs.

Chemotherapy, hormone therapy, and radiation therapy are currently used to treat women with bone metastatic BCa. While these treatments often shrink or slow the growth of bone metastases and can help alleviate symptoms associated with bone metastasis, they usually do not eliminate the metastases completely. Targeted antibody therapies, including trastuzumab and pertuzumab, are established standards of care for HER2-positive adjuvant and metastatic BCa. However, the poor bioavailability of these agents within bone tissues have limited their efficacy in dealing with HER2-positive bone metastases,^19–22^ In a recent long-term follow-up study of patients with HER2-positive metastatic breast cancers who received chemotherapy and trastuzumab, only 17% of bone metastatic BCa patients experienced a complete response, and none experienced a durable complete response. By comparison, a 40% complete response and 30% durable complete response was achieved in BCa patients with liver metastases.^22^ Thus, therapies with improved outcomes for BCa patients with bone metastases are highly desired.

In this study, we have used conjugation of bone-targeting moieties to develop an innovative bone targeting (BonTarg) technology that enables the preparation of antibodies with both antigen and bone specificity. Our data suggest that modification of the therapeutic HER2 antibody trastuzumab (Tras) with the bone-targeting bisphosphonate molecule, Alendronate (ALN), results in enhanced conjugate localization within the bone metastatic niche, relative to other tissues, raising the intriguing possibility that the bone-targeting antibody represents an enhanced targeted therapy for patients with bone metastases. We have tested this hypothesis using two breast cancer BMM models. The bone-targeting antibody conjugate, Tras-ALN, retains all the mechanistic properties of unmodified Tras, but exhibits enhanced ability to inhibit further BCa metastasis within the bone as well as metastasis-to-metastasis seeding from bone lesions. We find that, compared to either ALN or Tras separately, the Tras-ALN conjugate represents a superior treatment for HER2-positive tumor cell-derived BMMs. Interestingly, BMMs in BCa patients with HER2-negative tumors can actually express HER2 and may rely on HER2 signaling for progression.^48,49^ Similarly, we also find that Tras-ALN is effective in treating BMMs in a model of HER2-negative bone metastasis, providing a new therapeutic strategy using ALN-Tras to reduce latent metastases that occur in some HER2-negative breast cancer patients. The affinity of ALN for bone tissue helps overcome physical and biological barriers in the bone microenvironment that impede delivery of therapeutic antibodies, thereby enriching and retaining Tras in the bone. The Tras-ALN conjugate may reach higher concentrations in the bone metastatic niche, relative to healthy bone tissues, due to the low pH of bone tumor sites.^50^ This is consistent with previous observations that bisphosphonate molecules prefer to bind to the bone matrix in an acidic tumor environment.^32–35^

The evolution of current antibody therapy has been focused on targeting new biomarkers and functionalizing it with novel cytotoxic payloads. In this study, we explore the potential benefits of adding tissue specificity to antibody therapy. Using novel BonTarg technology, we have prepared the first bone-targeting antibodies by site-specific modifying with bone-targeting moieties. The resulting bone-targeting antibodies exhibit improved *in vivo* therapeutic efficacy in the treatment of breast cancer micrometastasis and in the prevention of secondary metastatic dissemination from the initial bone lesions. This type of precision delivery of biological medicines to the bone niche represents a promising avenue for treating bone-related diseases. The enhanced therapeutic profile of our bone-targeted HER2 antibody in treating microscopic BCa bone metastases will inform the potential benefit of adding tissue-specificity to traditional therapeutic antibodies.

## MATERIALS AND METHODS

### Construction of Tras-ALN conjugates

The non-canonical amino acid azide-Lys was incorporated at the C terminus of the ssFB-FPheK peptide via solid-phase peptide synthesis (Fig. S2). After HPLC purification, the peptide was denatured with 6 M urea and stepwise dialyzed to remove urea and allow peptide refolding. After buffer-exchange into PBS (pH 8.5), 32 equiv of ssFB-azide peptide was co-incubated with Tras (BS046D from Syd labs) in PBS (pH 8.5) buffer at 37 °C for two days. The Tras-azide conjugate was then purified via a PD-10 desalting column to remove excess ssFB-azide. The Tras-azide conjugate was characterized by ESI-MS. ESI-MS: expect 53564, found: 53558 (Fig. S3). 10 equiv of BCN-ALN was added to the solution at RT over night to selectively react with the azide group on the conjugate. Finally, the ALN labelled antibody conjugate was purified via a PD-10 desalting column to remove excess ALN-BCN. The conjugate was characterized by ESI-MS. ESI-MS: expect 53988, found: 53984 (Fig. 1C).

### Cell lines

MDA-MB-361, MCF-7, BT474, SK-BR-3, and MDA-MB-468 cell lines were cultured according to ATCC instructions. Firefly luciferase and GFP labelled MDA-MB-361 and MCF 7 cell lines were generated as previously described.^51^

### HA binding assay

Briefly, Tras or ALN-Tras was diluted in 1 mL PBS in an Eppendorf tube. Hydroxyapatite (15 equiv, 15 mg) was added, and the resulting suspension was shaken at 220 rpm at 37°C. Samples without hydroxyapatite were used as controls. After 0.25, 0.5, 1, 2, 4 and 8 hours, the suspension was centrifuged (3000 rpm, 3 min) and the absorbance of the supernatant at 280 nm was measured by Nanodrop. The percent binding to HA was calculated as follow, where OD represents optical density:

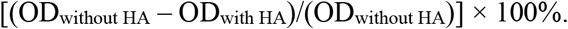

### Native bone binding assay

Long bones of mice were cut into small fragments, washed with distilled H2O and anhydrous ethanol, and then dried at room temperature overnight. For binding studies, Tras or ALN-Tras was diluted in 1 mL PBS in an Eppendorf tube. 30 mg dried bone fragments were added to the tube, and the resulting suspension was shaken at 220 rpm at 37°C. Samples without bone fragments were used as controls. After 0.25, 0.5, 1.0, 2.0, 4.0 and 8.0 h, the suspensions were centrifuged (3000 rpm, 5 min) and the absorbance at 280 nm of the supernatant was measured by Nanodrop. The percent binding to native bone was calculated according to the following formula, where OD represents optical density:

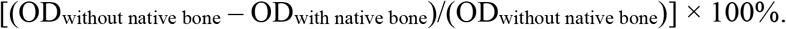

### *In vitro* cytotoxicity of Tras and Tras-ALN

SK-BR-3, BT474, and MDA-MB-468 cells were seeded in 200 *μ*L of culture medium into 96-well plates at a density of 2 × 10^3^ cells/well and incubated overnight to allow attachment. Culture medium was then removed, replaced by different concentrations of Tras and Tras-ALN dissolved in culture medium, and then incubated for 4 d. 20 *μ*L of MTT solution (5 mg/mL) was then added to each well and incubated for another 4 h. Medium was aspirated and 150 *μ*L DMSO was added to each well. The absorbance at 570 nm was measured by microplate reader (Infinite M Plex by Tecan) to quantify living cells.

### Flow cytometry

Cancer cells (3 × 10^3^) were re-suspended in 96-well plates and stained with 30 μg/mL Tras and Tras-ALN for 30 min at 4°C. After staining, cells were washed twice with PBS and then further incubated with Fluorescein (FITC) AffiniPure Goat Anti-Human IgG (H+L) (code: 109-095-003, Jackson Immunology) for 30 min at 4°C. Fluorescence intensity was determined using a BD FACSVerse (BD Biosciences).

### Determination of Kd values

The functional affinity of Tras-ALN for HER2 was determined as reported.^52^ Briefly, 2 × 10^5^ SK-BR-3, BT474, MDA-MB-361, or MDA-MB-468 cells were incubated with increasing concentrations of Tra and Tras-ALN for 4 hours on ice. After washing away unbound material, bound antibody was detected using Fluorescein (FITC) AffiniPure Goat Anti-Human IgG (H+L) (Jackson Immunology). Cells were analyzed for fluorescence intensity after propidium iodide (Molecular Probes, Eugene, OR) staining. The linear portion of the saturation curve was used to calculate the dissociation constant, KD, using the Lineweaver-Burk method of plotting the inverse of the median fluorescence as a function of the inverse of the antibody concentration. The KD was determined as follows: 1/F=1/Fmax+(KD/Fmax)(1/[Ab]), where F corresponds to the background subtracted median fluorescence and Fmax was calculated from the plot.

### Confocal imaging

Cancer cells were grown to about 80% confluency in 8-well confocal imaging chamber plates. Cells were incubated with 30 nM Tras-FITC for 30 min and then fixed by 4% paraformaldehyde for 15 min. Cells were washed three times with PBS (pH 7.4) and then incubated with DiIC18(3) (Marker Gene Technologies, Inc.) for 20 min and Hoechst 33342 (Cat No: H1399, Life Technologies™.) for 5 min. Cells were then washed three times with PBS (pH 7.4) and used for confocal imaging. Confocal fluorescence images of cells were obtained using a Nikon A1R-si Laser Scanning Confocal Microscope (Japan), equipped with lasers of 405/488/561/638 nm.

### Binding to bone cryosections

Nondecalcified long bone sections from C57BL/6 mice were incubated with 50 μg/mL Tras or Tras-ALN, conjugated overnight at 4 °C, followed by staining with fluorescein isothiocyanate (FITC)-labeled anti-human IgG for 60 min at room temperature. After washing 3 times with PBS, specimens were incubated for 30 min at 37 °C with Xylenol Orange (XO) (stock: 2 mg/ml, dilute 1:500, dilute buffer: PBS pH 6.5). After three washes with PBS, specimens were stained with Hoechst 33342 (stock 10mg/ml, dilute 1:2000) for 10 min. Slides were then washed with PBS, air dried, and sealed with Prolong™ gold anti-fade mountant (from ThermoFisher).

### *In vivo* evaluation of Tras-ALN

Intra-iliac injections and IVIS imaging were performed as previously described.^53^ Five days after injection, animals were randomized into four groups: PBS treated control, ALN (a representative of free BP, retro-orbital injection 10 μg/kg in PBS twice a week), Tras (1.0 mg/kg retro-orbital injection in sterile PBS twice a week), and Tras-ALN conjugate (same as Tras). After injection, animals were imaged twice a week using IVIS Lumina II (Advanced Molecular Vision), following the recommended procedures and manufacturer’s settings. On day 110, mice were anesthetized and blood was collected by cardiac puncture prior to euthanasia. Tumor-bearing tibia, heart, liver, spleen, lung, brain and kidney were collected for further tests.

### *Ex vivo* metastasis-to-metastasis analysis

Mice were anesthetized with 2.5 % isoflurane in oxygen and injected with luciferin retro-orbitally. Mice were then euthanized and their hearts, livers, spleens, lungs, kidneys, brain, and tibia bones were collected. *Ex vivo* bioluminescence and fluorescence imaging of these organs were immediately performed on the IVIS Lumina.

### Bone histology and immunohistochemistry

Harvested long bones were fixed for 1 week in 10% formalin and then decalcified in 12% EDTA at 4°C for two weeks. Specimens were embedded in paraffin using the standard procedure. From these blocks, 5 μm sections were cut and collected on glass slides. Sections were dried in an oven overnight (37°C) and then deparaffinized in xylene solution for 10 min. Hematoxylin and eosin (H&E) staining were performed via the conventional method. Immunohistochemistry analysis was performed on decalcified paraffin-embedded tissue sections using the HRP/DAB ABC IHC KIT (abcam) following the manufacture’s protocol.

### Radiographic analysis

Tibiae were dissected, fixed and scanned by microcomputed tomography (micro-CT, Skyscan 1272, Aartselaar, Belgium) at a resolution of 6.64 μm/pixel. Raw images were reconstructed in NReconn and analyzed in CTan (SkyScan, Aartselaar, Belgium) using a region of interest (ROI). Bone parameters analyzed included trabecular thickness (Tb.Th), bone volume fraction (BV/TV), bone mineral density (BMD), and BS/BV (bone surface/bone volume ratio).

### Biodistribution

MDA-MB-361 cells were introduced into female athymic nude mice (body weight = 13-15 g) via intra-iliac injections. After three months, Cy7.5-labeled Tras and Tras-ALN (1 mg/kg) were administrated to tumor-bearing nude mice by retro-orbital injection. At 24 h, 96 h, or 168 h after injection, major organs including heart, liver, spleen, kidney, lung, and bone tumor tissue were removed. The fluorescence intensity in organs and bone tumor tissues was determined semiquantitatively by using the Caliper IVIS Lumina *in vivo* imager (Caliper Life Science, Boston, MA, USA). Bones from Tras-ALN treated mice were fixed and sectioned to further evaluate bio-distribution.

In a separate study, unlabeled Tras-ALN (1 mg/kg) was administered via retro-orbital injection to nude mice bearing MDA-MB-361 tumors in their right hind limbs. After 48 hours, long bones from Tras-ALN treated mice were isolated and immediately sectioned without decalcification. Bone sections were then fixed and incubated with anti-RFP (rabbit) antibody (1:200, purchased from Rockland) overnight at 4, followed by staining with fluorescein isothiocyanate (FITC)-labeled anti-human IgG (1:100, purchased from Jackson Immunology) and Alexa Fluor® 555 AffiniPure Donkey Anti-Rabbit IgG (H+L) (1:200, purchased from Thermo Fisher) for 120 min at room temperature. Sections were mounted with Prolong™ gold anti-fade mountant with DAPI (from ThermoFisher) and sealed with a coverslip, then used for confocal imaging.

### Quantification of TRAP and calcium levels in serum

At terminal time points, blood was collected by cardiac puncture, and centrifuged for 15 min at 3,000 rpm to obtain the serum. The concentration of osteoclast-derived TRACP 5b was measured by using a Mouse ACP5/TRAP ELISA Kit (catalog number IT5180, GBiosciences). Serum calcium levels were determined colorimetrically using a calcium detection kit (catalog number DICA-500, Bioassays).

### Statistical methods

Data are presented as means plus or minus SEM and statistically analyzed using GraphPad Prism software version 6 (GraphPad software, San Diego, CA). Two-way ANOVA followed by Sidak’s multiple comparisons was used for all data collected over a time course. One-way ANOVA followed by Tukey’s multiple comparisons was used for Micro-CT data. Unpaired Student’s *t*-test was used for multi-organ metastasis data. P < 0.05 was considered to represent statistical significance.

## Acknowledgments

We thank Drs. Xiao and Zhang Laboratory members for insightful comments. This work was supported by the Cancer Prevention Research Institute of Texas (CPRIT RR170014 to H.X.), NIH (R35-GM133706 to H.X., NCI CA183878, and NCI CA221946 to X.H.-F.Z.), the Robert A. Welch Foundation (C-1970 to H.X.), US Department of Defense (DAMD W81XWH-16-1-0073, Era of Hope Scholarship to X.H.-F.Z.), the John S. Dunn Foundation Collaborative Research Award (to H.X.), and the Hamill Innovation Award (to H.X.). H.X. is a Cancer Prevention & Research Institute of Texas (CPRIT) scholar in cancer research. X.H.-F.Z. is also supported by the Breast Cancer Research Foundation and McNair Medical Institute.

## Author contributions

Z.T., L.W., X.H.-F.Z. and H.X. developed the hypothesis, designed experiments, analyzed the data, and wrote the manuscript. Z.T., L.W., C.Y., Z.X., I.B., H.W., and W.Z. performed experiments. Y.C., A.L., L.W., K.W., X.H.-F.Z. and H.X. contributed to experimental design, generation of the reagents, and manuscript editing. X.H.-F.Z. and H.X. conceived and supervised the project.

