## Supporting Information for "Harnessing the Power of Antibodies to Fight Bone Metastasis"

**Synthesis of BCN-ALN.** BCN-PNP (ENDO) (31.5 mg) and DIPEA (38.7 mg) were dissolved in 1 mL dimethylsulfoxide (DMSO), followed by dropwise addition of 27.4 mg of ALN (dissolved in 0.3 mL of deionized water) into the mixture. The resulting mixture was stirred for 4 hours. Ethyl acetate (1 mL) was added to the reaction solution, and the resulting precipitate was filtered and rinsed three times with ethyl acetate. The product was purified by reversed-phase column chromatography. The structure of BCN-ALN was confirmed by MS. ESI-MS $[M-H]^-$: Calcd. For $C_{15}H_{25}NO_9P_2$ 424.1, found: 424.1 (Fig. S1).

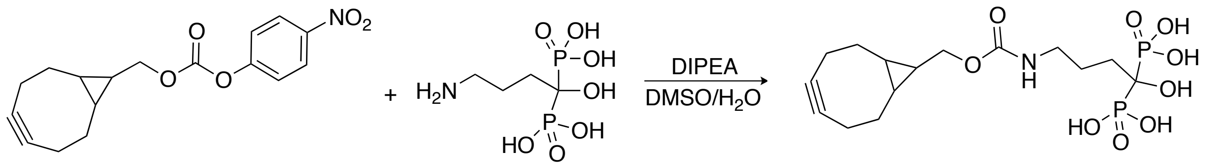

**Solid-phase synthesis of ssFB-FPheK peptide.** ssFB peptide was synthesized by following the protocol for Fmoc-based peptide synthesis. Amino acid sequence: FNKEQQNAFYEILHLPNLNXEQRNAFIQSLKDDPS-AzK (X=MMT-Lys). Rink Amide MBHA resin was used as the solid support. The Fmoc protection group was removed using 25% piperidine in DMF. After washing the product five times with DMF, pre-activated HATU/Fmoc-amino acid mixture in DMF was added to the reaction vessel for amide bond formation. The next amino acid was then coupled to the beads via the same reaction cycle. Once peptide synthesis was complete, the N-terminus was capped by acetic anhydride. In order to incorporate FPheK into the peptide, the MMT protection group was first selectively removed using 10% acetic acid (AcOH:TFE:DCM=1:2:7). Subsequently, 2 equiv of 4-fluorophenyl chloroformate and 4 equiv of DIEA were added into the reaction vessel to react with the exposed free amine on the Lys side chain for FPheK formation. Once the reaction was complete, appropriate quantities of TFA and scavengers (water, anisole, triisopropyl silane) were added to the vessel to cleave the peptide from the resin, and to remove and quench all protection groups. The peptide was then precipitated by addition of ice-cold ether, purified by HPLC and lyophilized. ESI-MS $[M+H]^+$: Calcd. For 4524, found: 4524 (Fig. S2).

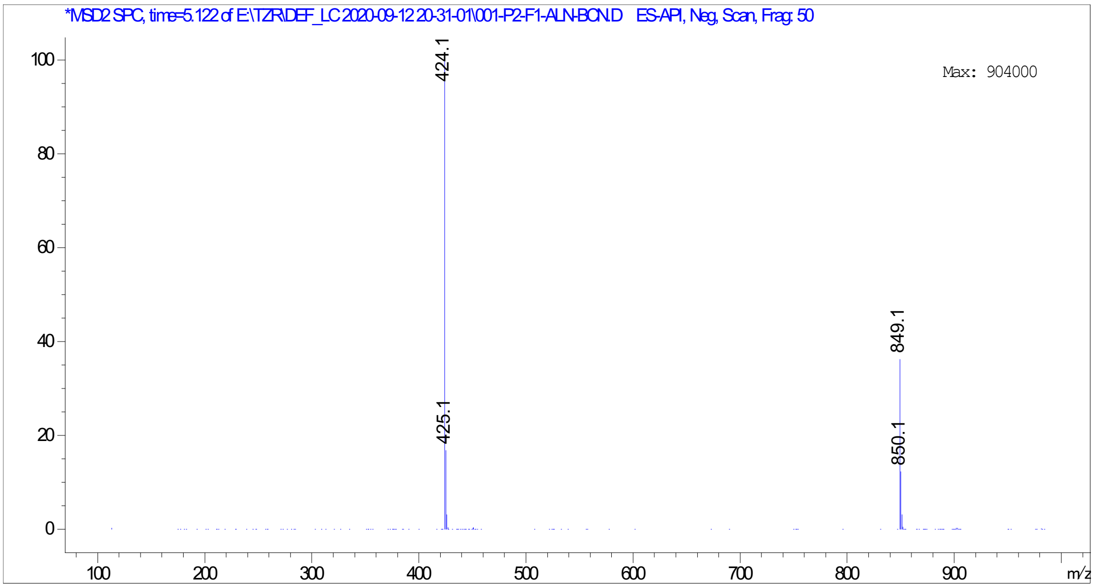

**Fig. S1 ESI-MS spectra of BCN-ALN.**

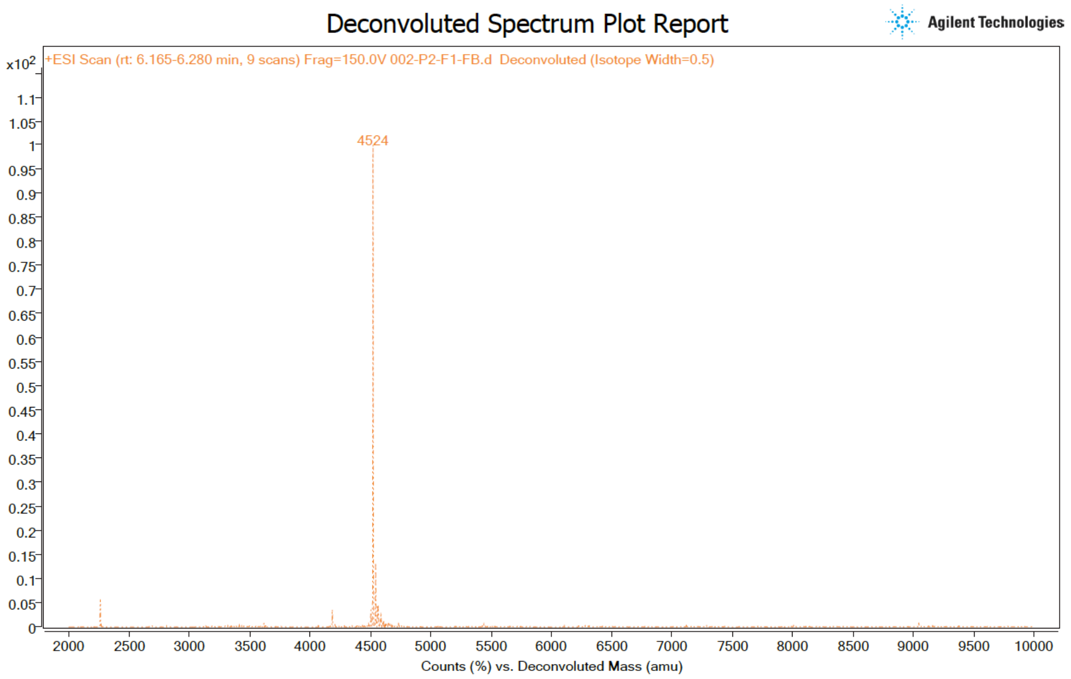

**Fig. S2 ESI-MS spectra of ssFB-FPheK**.

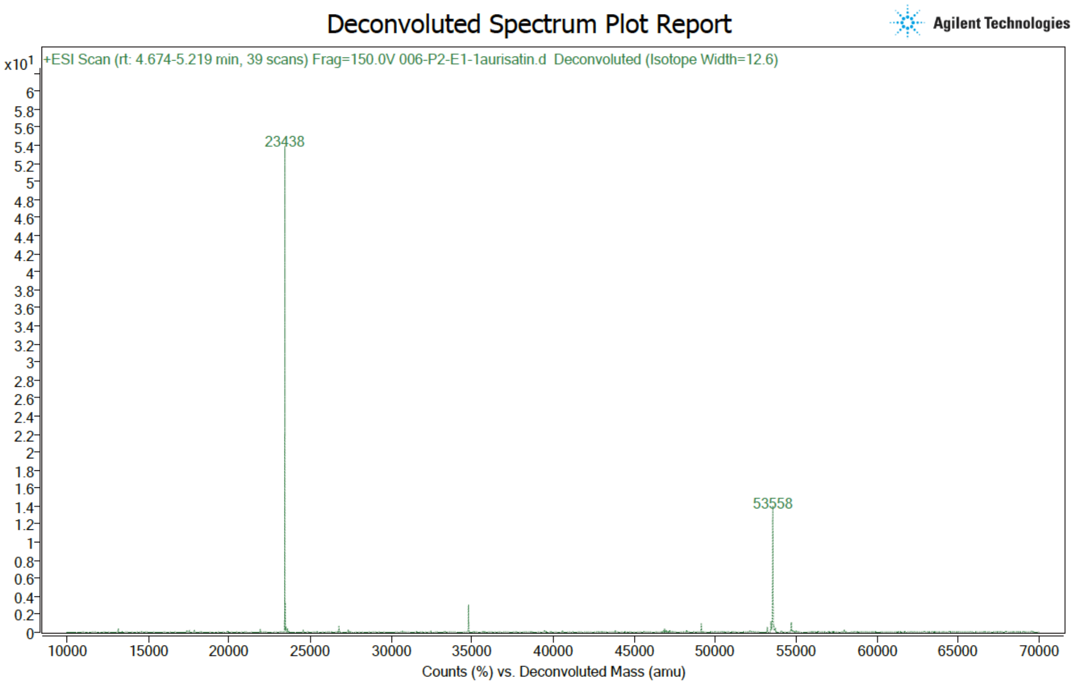

**Fig. S3 ESI-MS spectra of Tras-azide.**

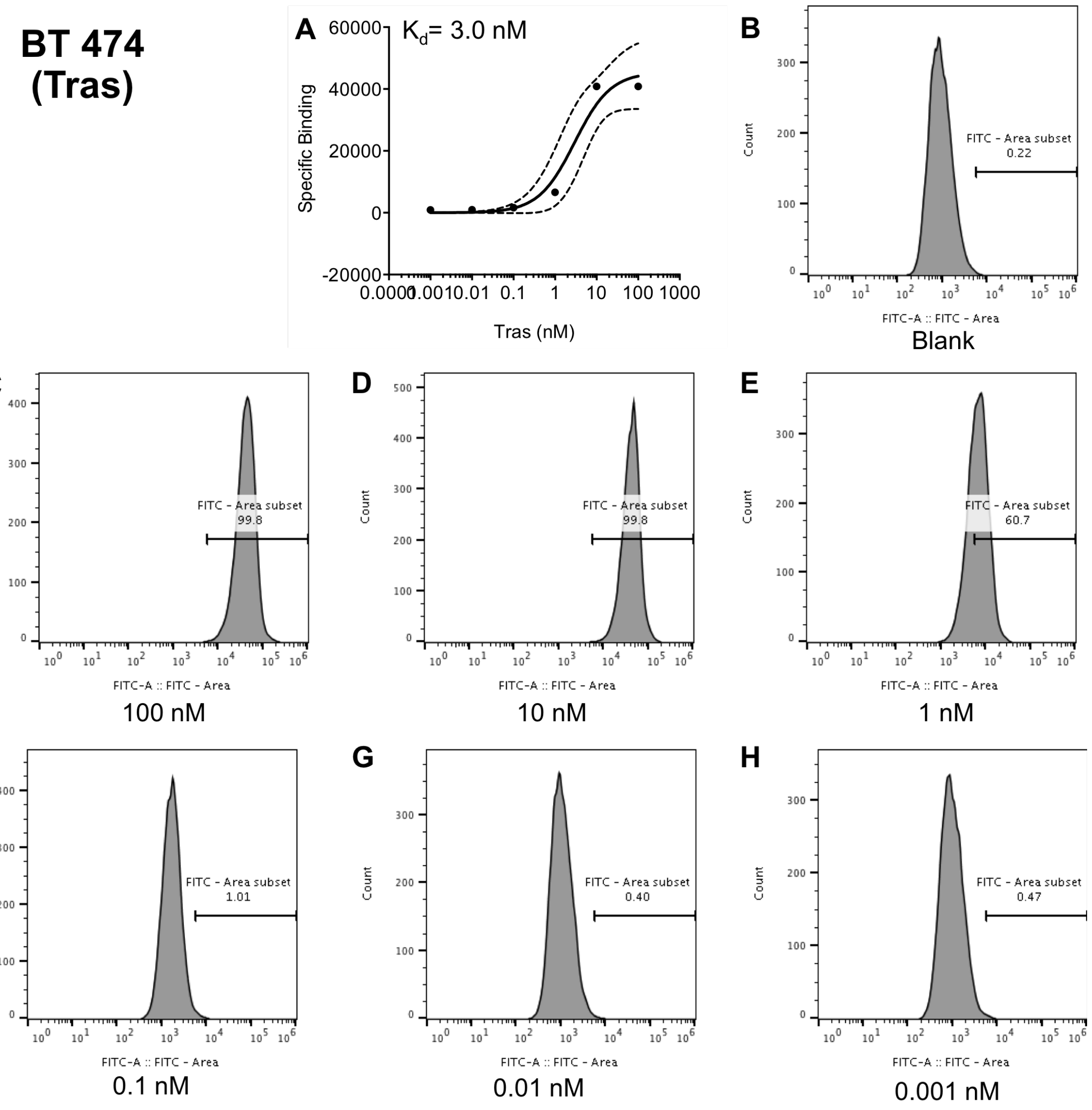

**Fig. S4 Tras binding to BT474 cells.** BT474 cells were incubated with increasing concentrations of Tras and process as described in Methods and fluorescence was measured on the flow cytometer. The KD was determined as follows: 1/F=1/Fmax+(KD/Fmax)(1/[Ab].

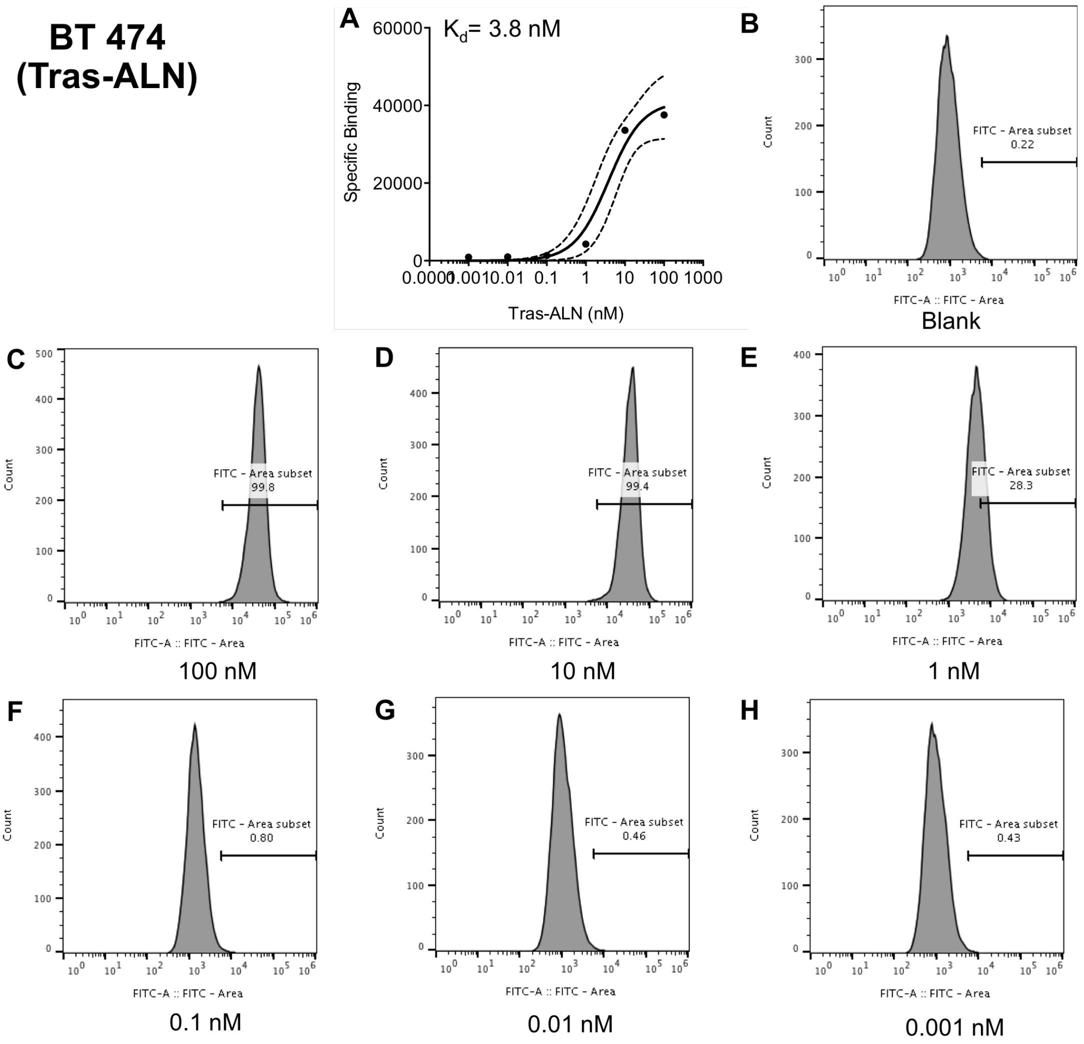

**Fig. S5 Tras-ALN binding to BT474 cells.** BT474 cells were incubated with increasing concentrations of Tras-ALN and process as described in Methods, and fluorescence was measured on the flow cytometer. The KD was determined as follows: 1/F=1/Fmax+(KD/Fmax)(1/[Ab].

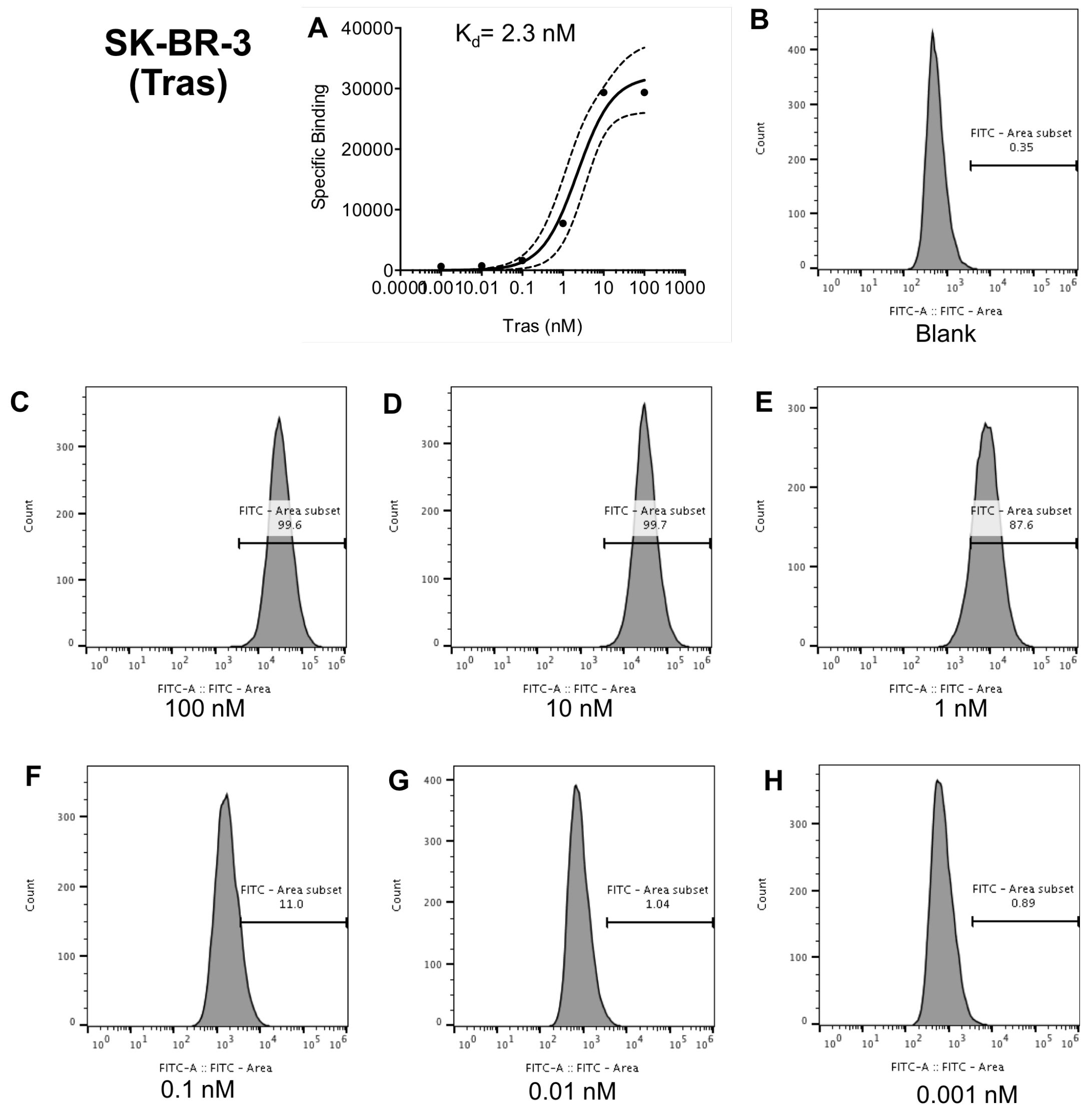

**Fig. S6 Tras binding to SK-BR-3 cells.** SK-BR-3 cells were incubated with increasing concentrations of Tras and process as described in Methods and fluorescence was measured on the flow cytometer. The KD was determined as follows: 1/F=1/Fmax+(KD/Fmax)(1/[Ab].

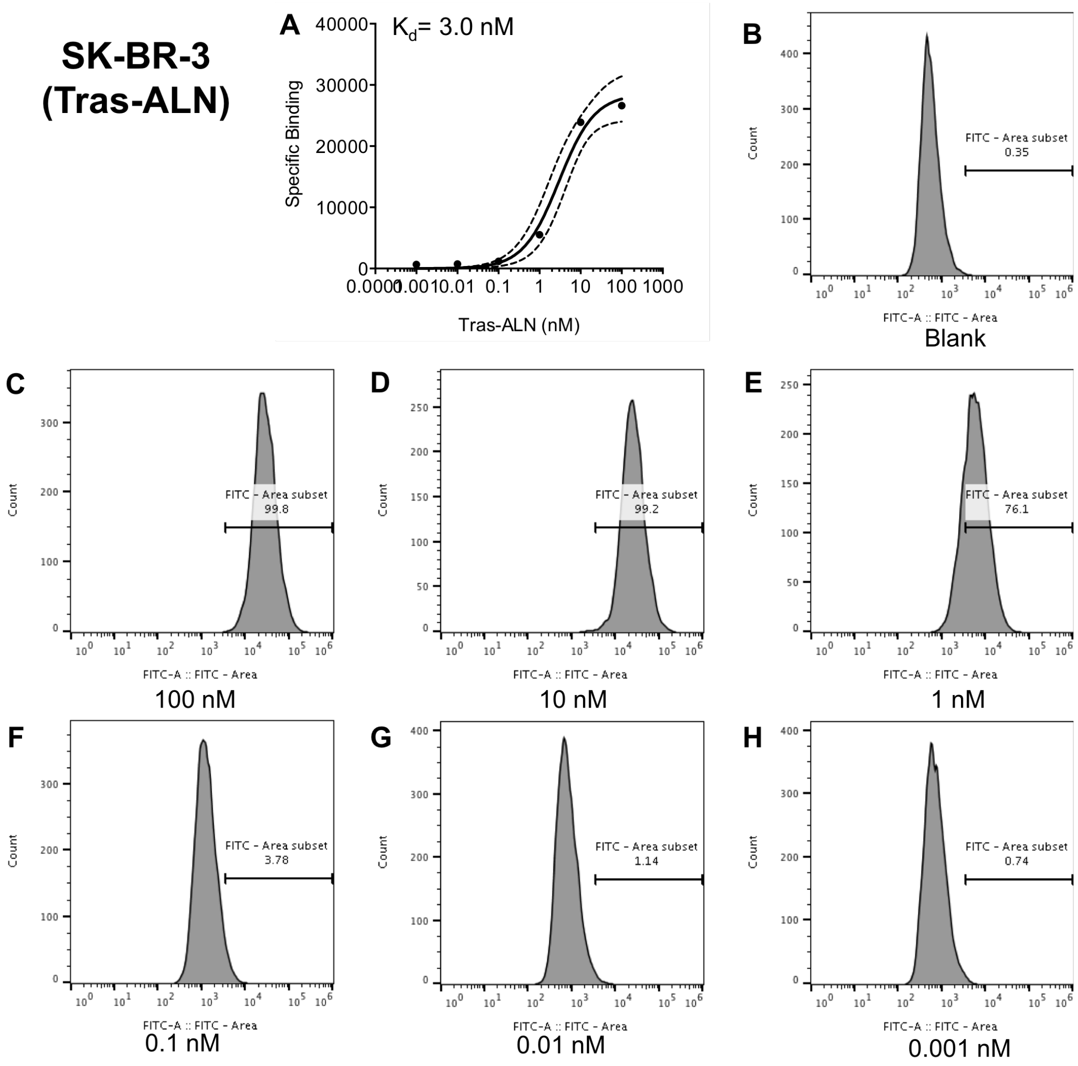

**Fig. S7 Tras-ALN binding to SK-BR-3 cells.** SK-BR-3 cells were incubated with increasing concentrations of Tras and process as described in Methods and fluorescence was measured on the flow cytometer. The KD was determined as follows: 1/F=1/Fmax +(KD/Fmax)(1/[Ab].

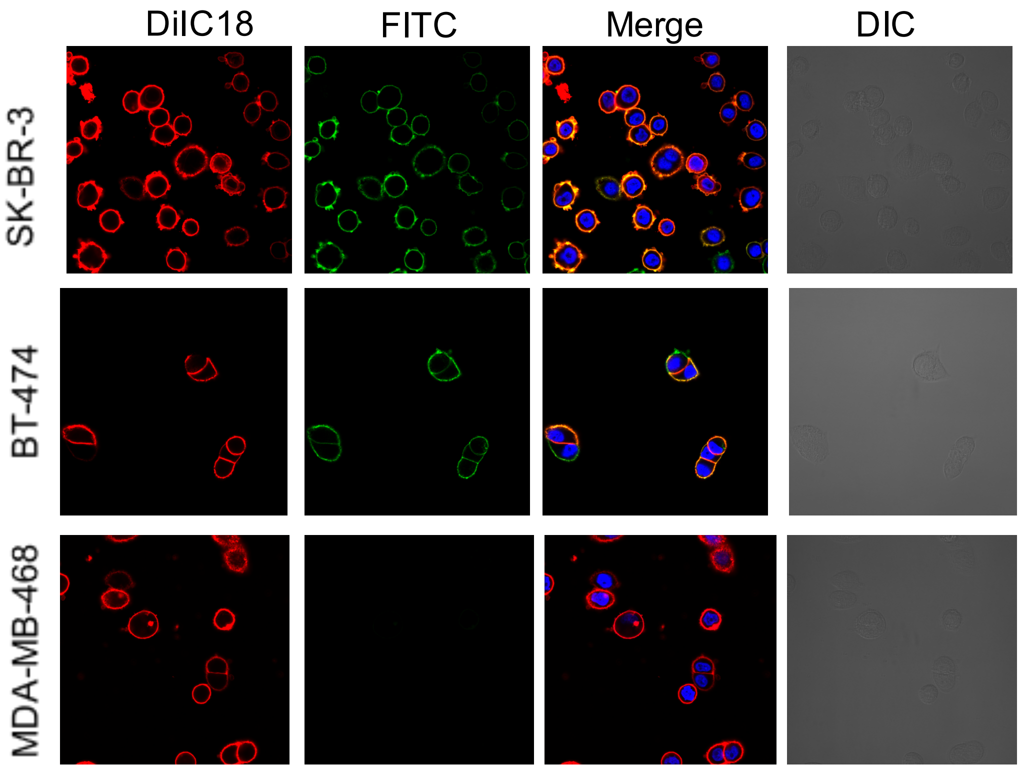

**Fig. S8 Binding of Tra-ALN in BT-474, SK-BR-3, and MDA-MB-468 cells visualized by confocal microscopy.** Cells were incubated with 30 nM Tras-ALN in media for 30 min at 37 °C and stained with DilC18 (red fluorescence) and Hoechst nuclear stain (blue fluorescence).

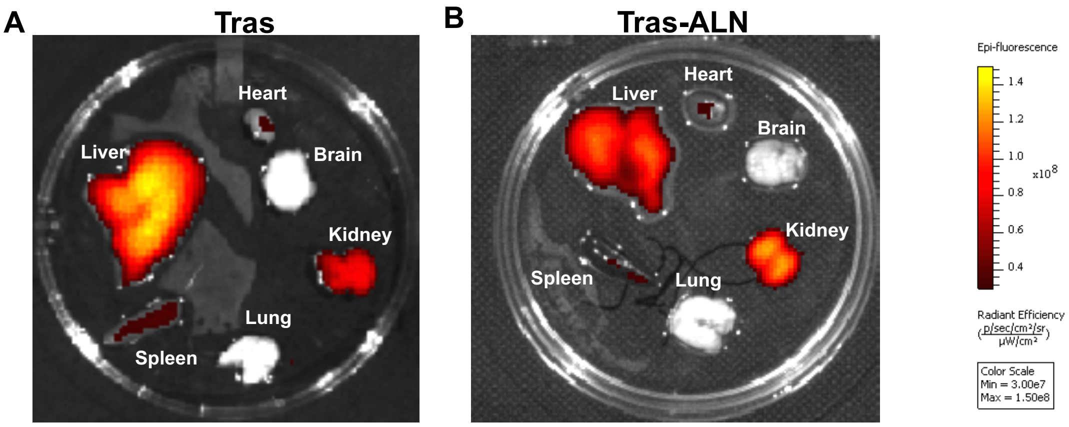

**Fig. S9** ***Ex vivo*** **fluorescence images of main organs.** Heart, liver, spleen, lung, kidney, brain of athymic nude mice bearing MDA-MB-361 tumors 96 h after the retro-orbital injection Cy7.5-labeled Tras and Tras-ALN.

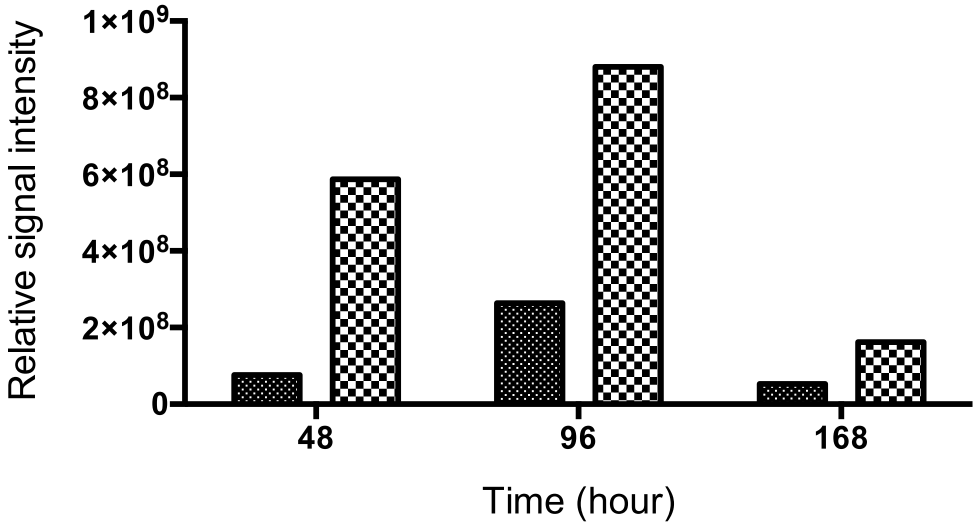

**Fig. S10 *Ex vivo* fluorescence images analysis for the bone biodistribution of Tras and Tras-ALN.** 24 h, 96 h or 168 h after after the retro-orbital injection of Cy7.5-labeled Tras and Tras-ALN. The bone was collected and analysis. The quantity data was summarized from the Fig. 1K. The signal of free tumor from Tras treated mice was considered as blank. The Relative signal was calculated as follows: The signal from hind limbs – The signal from free tumor hind limbs (from Tras treated group).

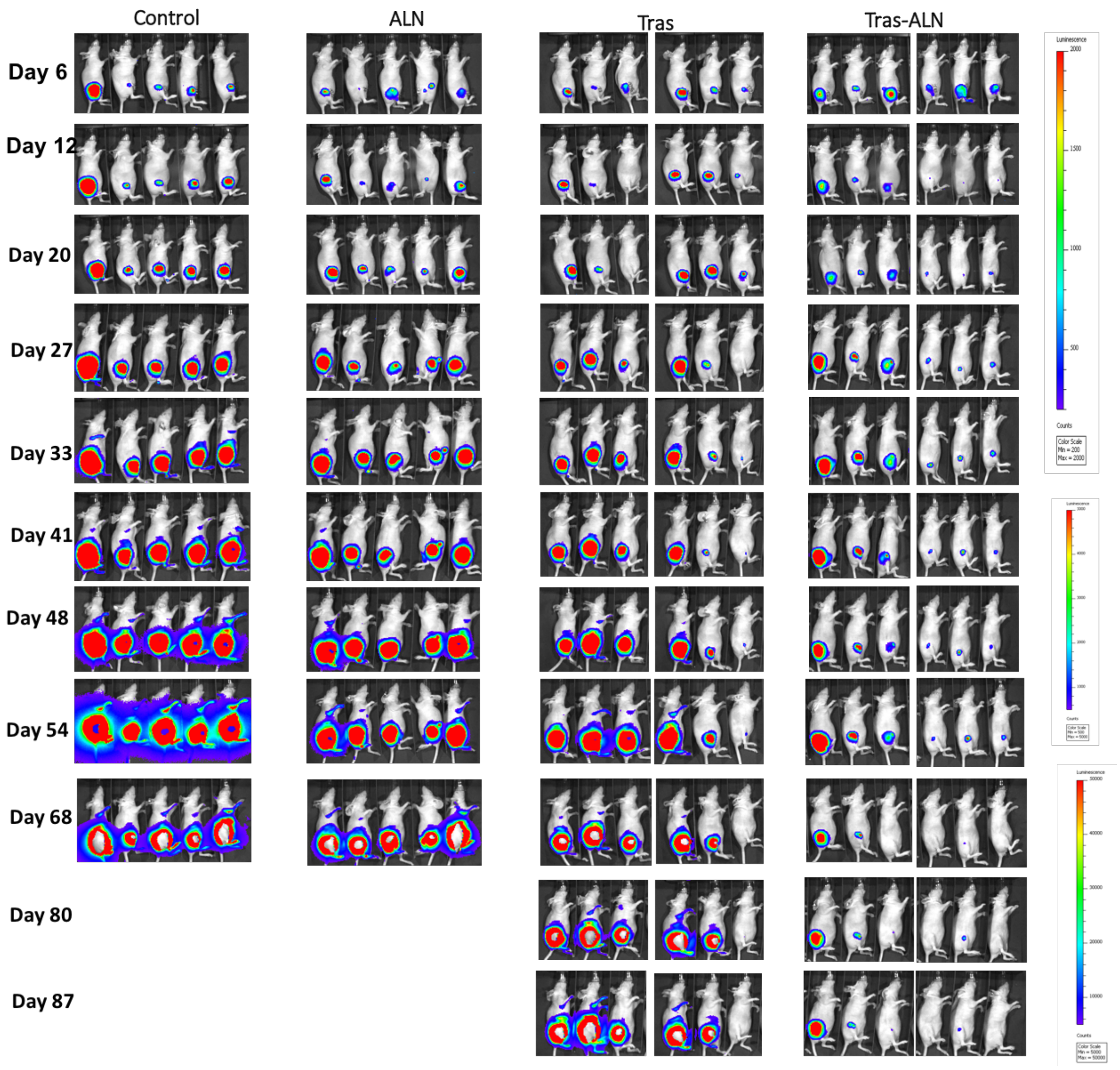

**Fig. S11 Tras-ALN inhibits breast cancer micrometastases in the bone.** MDA-MB-361 cells were IIA injected into the right hind limb of nude mice, followed by treatment with PBS, ALN (10 µg/kg retro-orbital venous sinus in PBS twice a week), Tras (1 mg/kg retro-orbital venous sinus in sterile PBS twice a week), and Tras-ALN conjugate (same as Tras). Tumor burden was monitored twice a week by bioluminescence imaging (Day 6, 20, 33, 48 and 68 imaging data were selected to show in Fig. 2A).

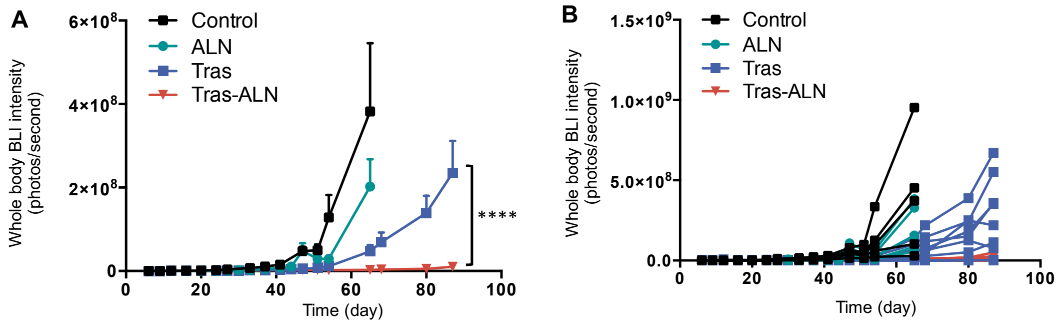

**Fig. S12 Whole body BLI quantification.** (A) The BLI from each treatment group quantified by the radiance detected in the whole body. (B) Individual whole body luminescent intensity of different treated group as described in Fig. 2A. ****$P < 0.0001$.

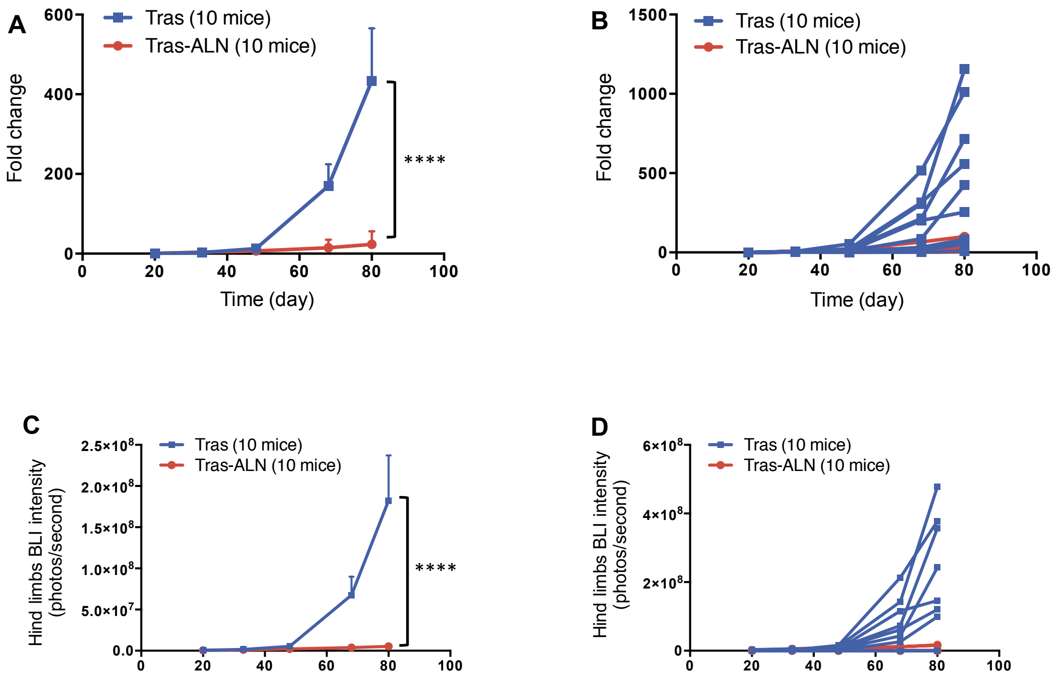

**Fig. S13 BLI signals in the hind limbs were quantified in Tras and Tras-ALN treated group and are shown.** (A) Fold-change in mean luminescent intensity of hind limbs in mice treated with Tras and Tras-ALN( as described in Fig. 2A), two-way ANOVA comparing Tras to Tras-ALN. (B) Fold-change in Individual luminescent intensity of hind limbs in Tras and Tras-ALN treated group. (C) The mean BLI of hind limbs from Tras and Tras-ALN treatment group quantified. (D) Individual luminescent intensity of Tras and Tras-ALN treated group. ****$P < 0.0001$.

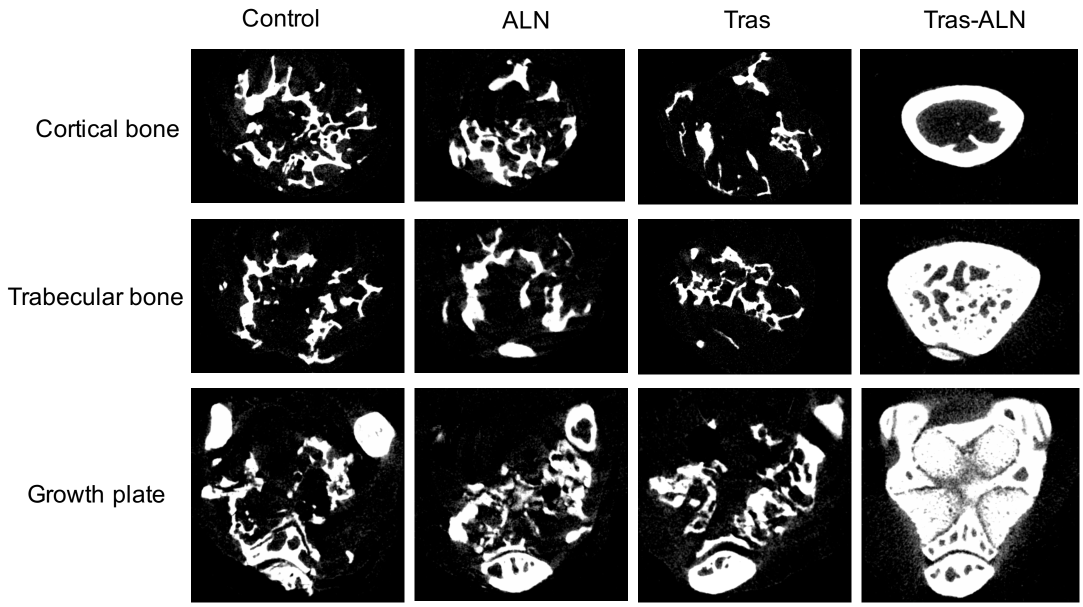

**Fig. S14 MicroCT-based 3D renderings of bones.** Cortical bone, images show extensive cortical bone destruction. Trabecular bone, images show trabecular destruction. Lower panel (growth plate), images plate show bone loss at growth plate.

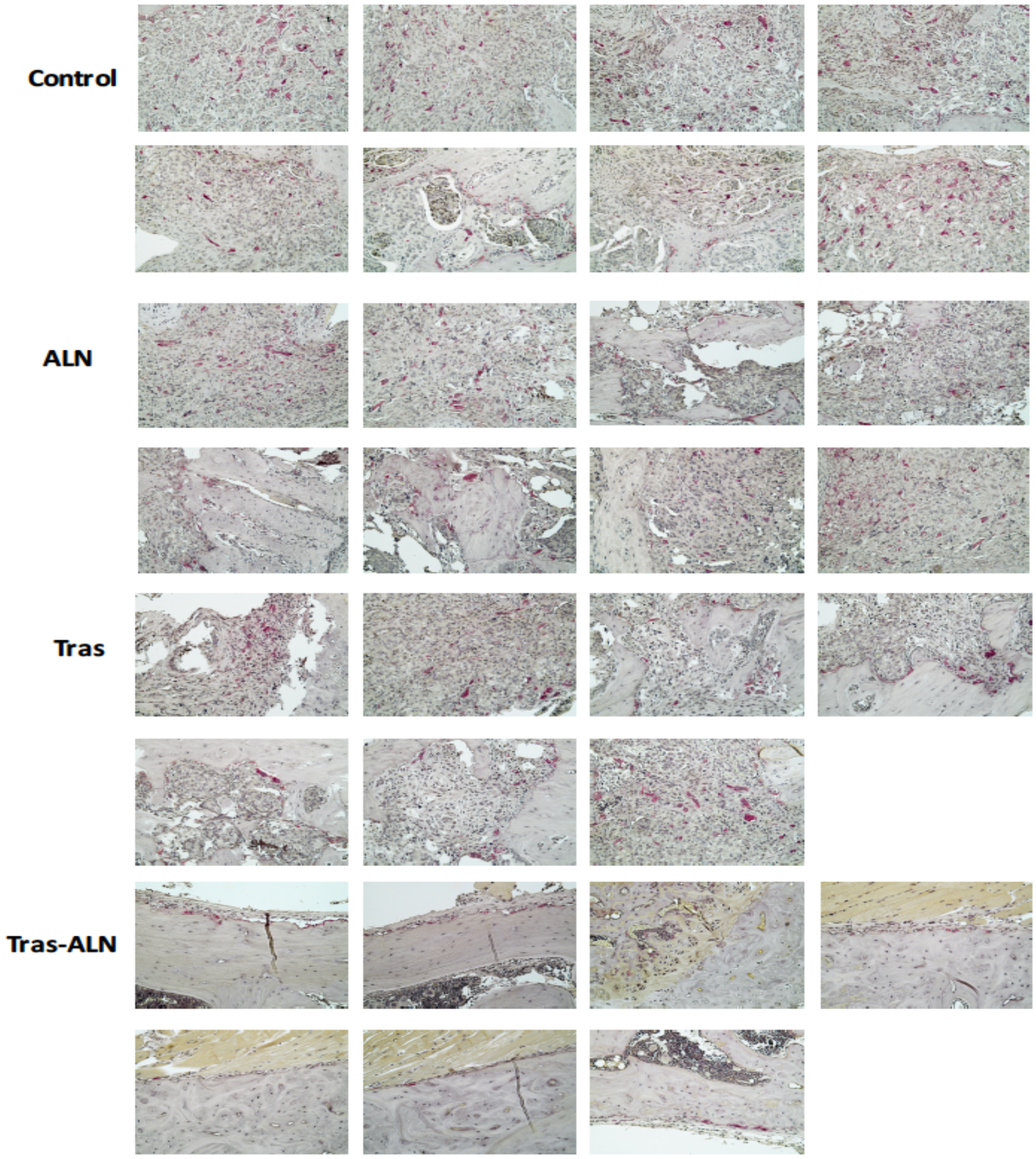

**Fig. S15 TRAP staining of bone sections from each group.**

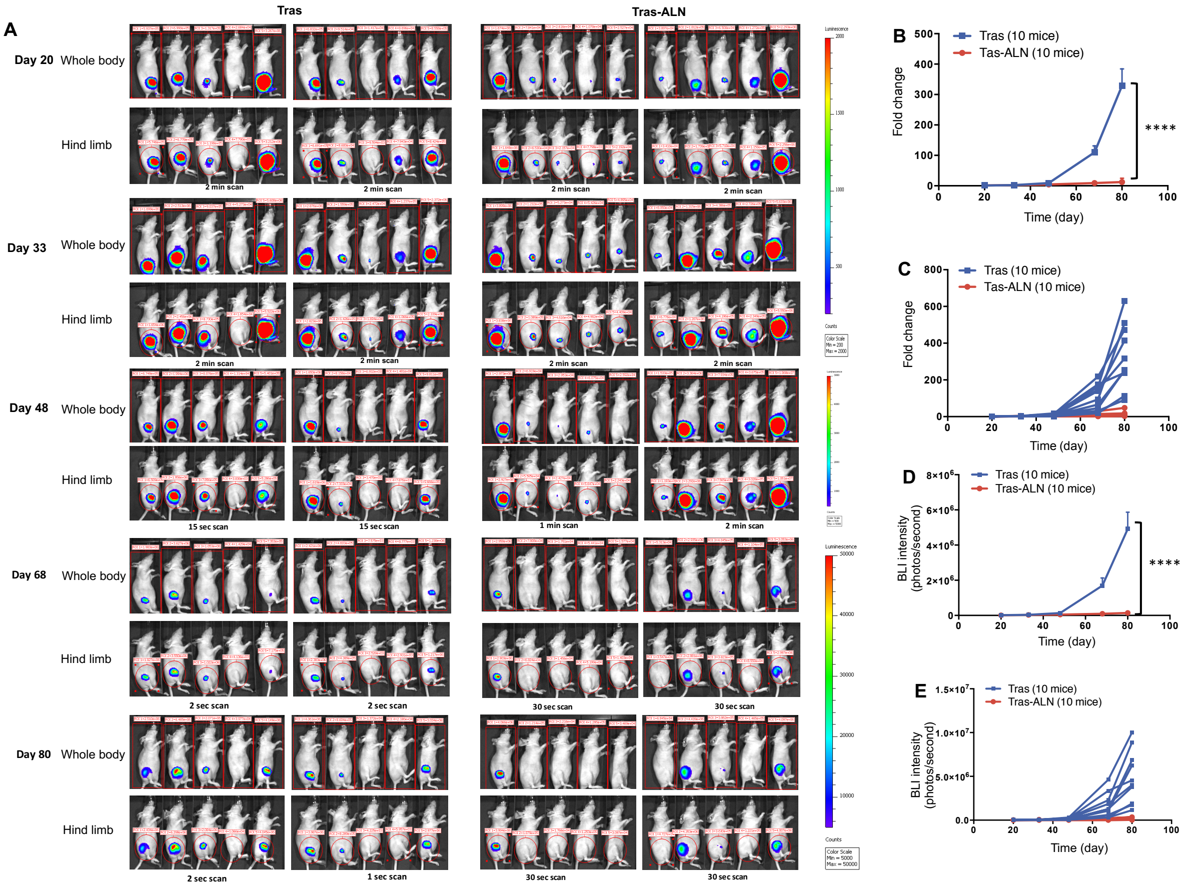

**Fig. S16 The *in vivo* quantification of secondary metastases.** (A) BLI signal in the whole body and hind limbs of mice in various treatment groups were quantified and are shown. The secondary metastases was determined as follows: , BLI signals in whole body and the hind limbs (shown by red circles) were quantified. Each time point, animals were imaged twice a week using IVIS Lumina II (Advanced Molecular Vision), following the recommended procedures and manufacturer's settings. For the groups which signal suggested "Saturated Luminescent Image", it will be scanned for shorter time (which were indicated under the imaging). The secondary metastases were calculated as follows: BLI signal intensity in whole body – BLI signal intensity in hind limbs. (B) Fold-change in mean luminescent intensity of secondary metastases in mice treated as described in (Fig. 2A), two-way ANOVA comparing Tras to Tras-ALN. (C) Fold-change in Individual luminescent intensity of secondary metastases in Tras and Tras-ALN treated group. (D) The mean BLI of secondary metastases from Tras and Tras-ALN treatment group quantified. (E) Individual luminescent intensity of Tras and Tras-ALN treated group. $****P < 0.0001$.

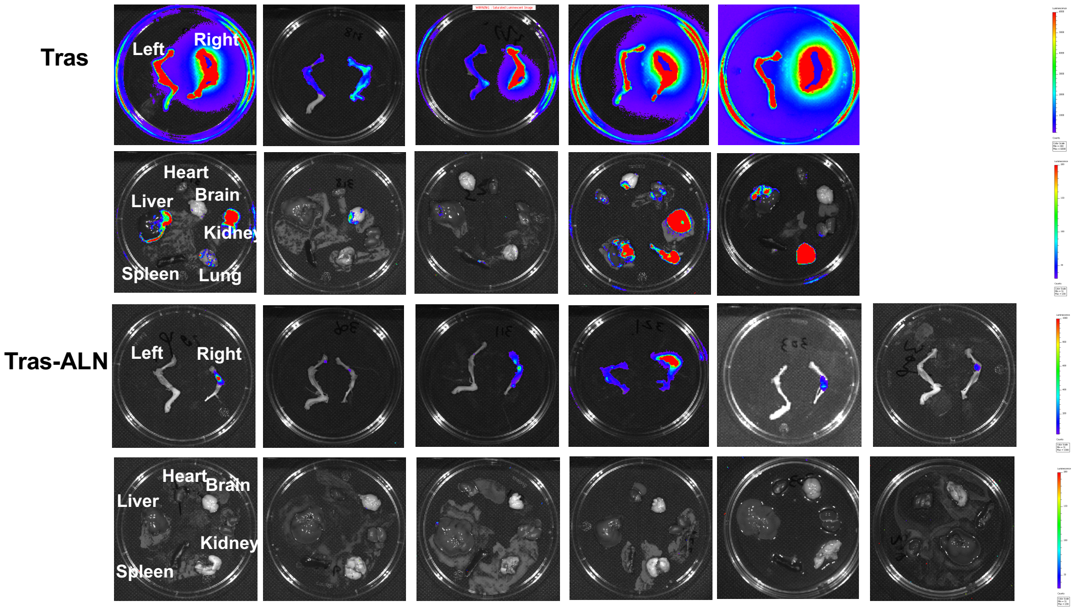

**Fig. S17 Secondary metastases observed in various organs in mice treated with Tras (top) or Tras-ALN (bottom).**

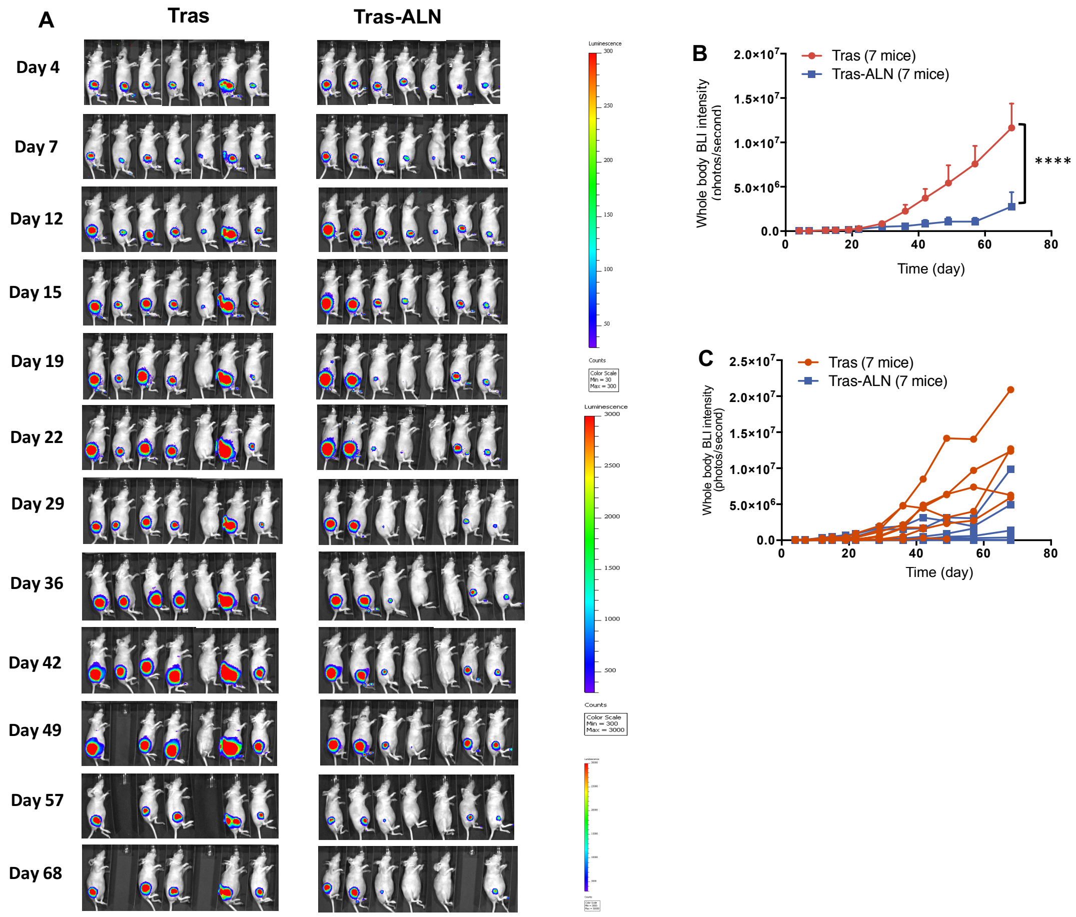

**Fig. S18 *In vivo* comparison of Tras and Tras-ALN in HER2-negative model.** MCF-7 cells were IIA injected into the right hind limb of nude mice, followed by treatment with Tras (1 mg/kg retro-orbital venous sinus in sterile PBS twice a week), and Tras-ALN conjugate (same as Tras). (A) Tumor burden was monitored twice a week by bioluminescence imaging (Day 4, 12, 19, 29 and 42 imaging data were selected to show in Fig. 4A. (B) The BLI from each treatment group quantified by the radiance detected in the whole body. (C) Individual whole body luminescent intensity of Tras and Tras-ALN treated group. ****$P < 0.0001$.

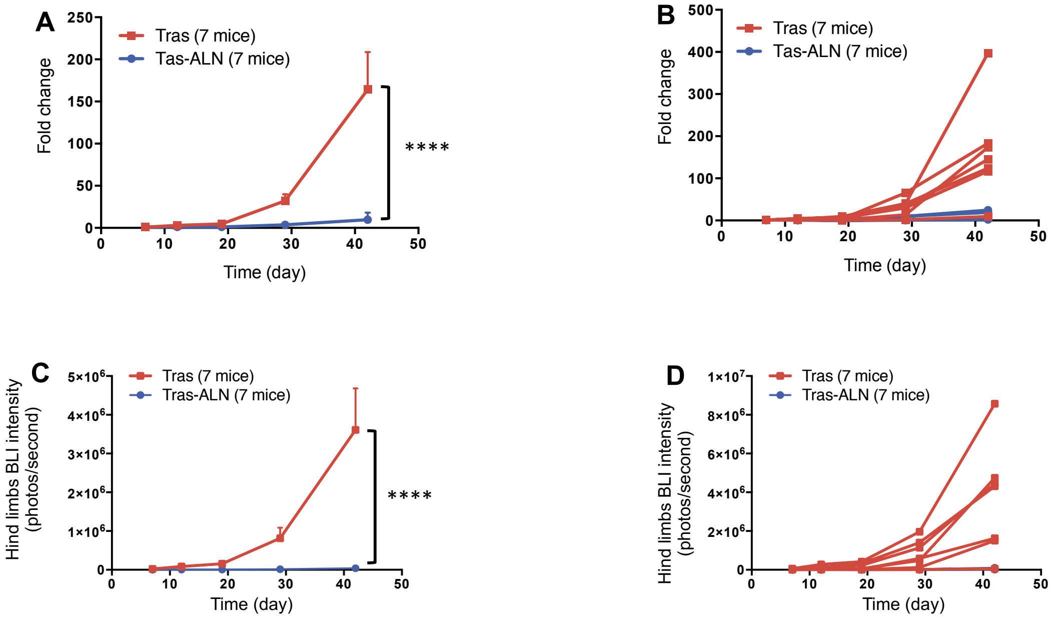

**Fig. S19** BLI signals in the hind limbs in MCF-7 model were quantified in Tras and Tras-ALN treated groups and are shown. (A) Fold-change in mean luminescent intensity of hind limbs in mice treated with Tras and Tras-ALN( as described in Fig. 4A), two-way ANOVA comparing Tras to Tras-ALN. (B) Fold-change in Individual luminescent intensity of hind limbs in Tras and Tras-ALN treated groups. (C) The mean BLI of hind limbs from Tras and Tras-ALN treatment groups quantified. (D) Individual luminescent intensity of Tras and Tras-ALN treated groups. ****$P < 0.0001$.

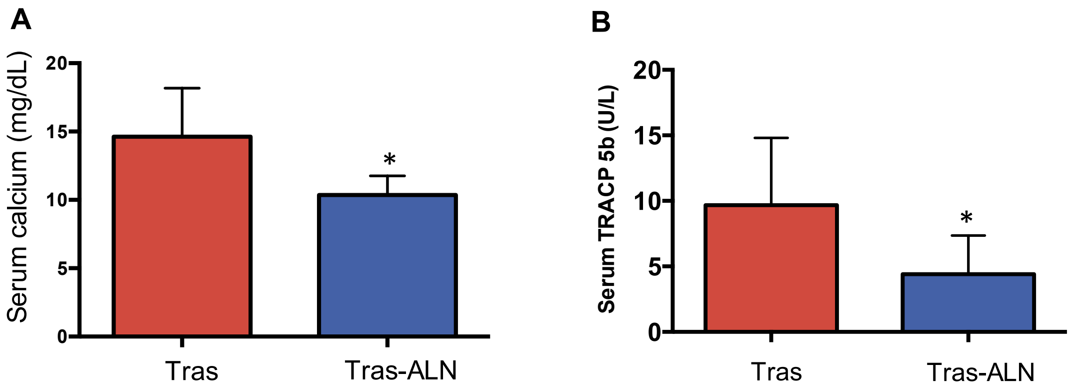

**Fig. S20** Effects of Tras-ALN on MCF-7 HER2-negative model: serum TRACP 5b and calcium levels analysis. (A) Serum TRACP 5b concentration in Tras and Tras-ALN at the end of experiment (*P < 0.05). (B) Serum calcium concentration in Tras and Tras-ALN group at the end of experiment(*$P$ < 0.05).

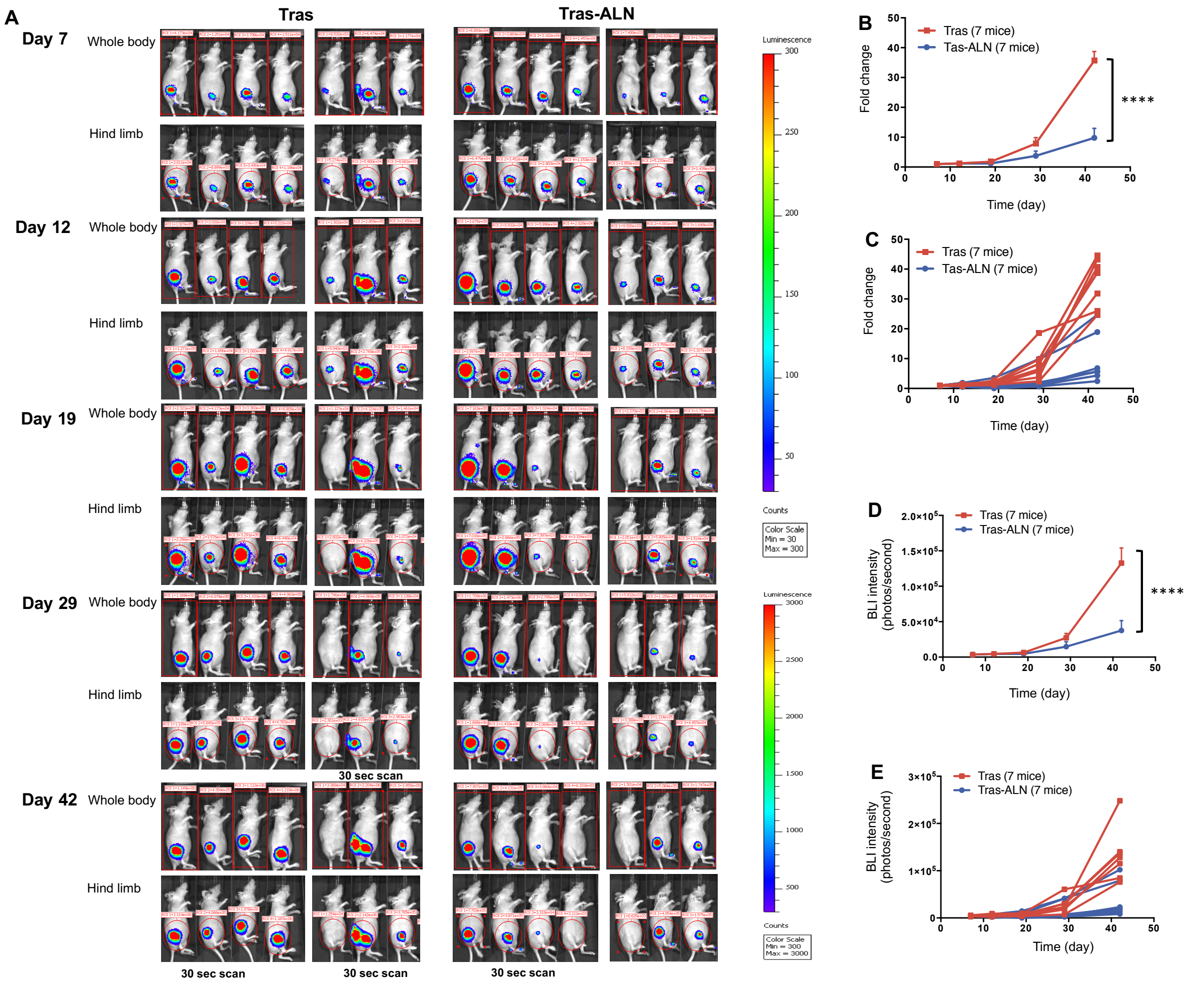

**Fig. S21 The *in vivo* quantification of secondary metastases.** (A) BLI signal in the whole body and hind limbs of mice in various treatment groups were quantified and are shown. The secondary metastases was determined as follows: , BLI signals in whole body and the hind limbs (shown by red circles) were quantified. Each time point, animals were imaged twice a week using IVIS Lumina II (Advanced Molecular Vision), following the recommended procedures and manufacturer's settings. For the groups which signal suggested "Saturated Luminescent Image", it will be scanned for shorter time (which were indicated under the imaging). The secondary metastases were calculated as follows: BLI signal intensity in whole body – BLI signal intensity in hind limbs. (B) Fold-change in mean luminescent intensity of secondary metastases in mice treated as described in Fig. S18, two-way ANOVA comparing Tras to Tras-ALN. (C) Fold-change in Individual luminescent intensity of secondary metastases in Tras and Tras-ALN treated groups. (D) The mean BLI of secondary metastases from each treatment group quantified. (E) Individual luminescent intensity of Tras and Tras-ALN treated groups. ****$P < 0.0001$.

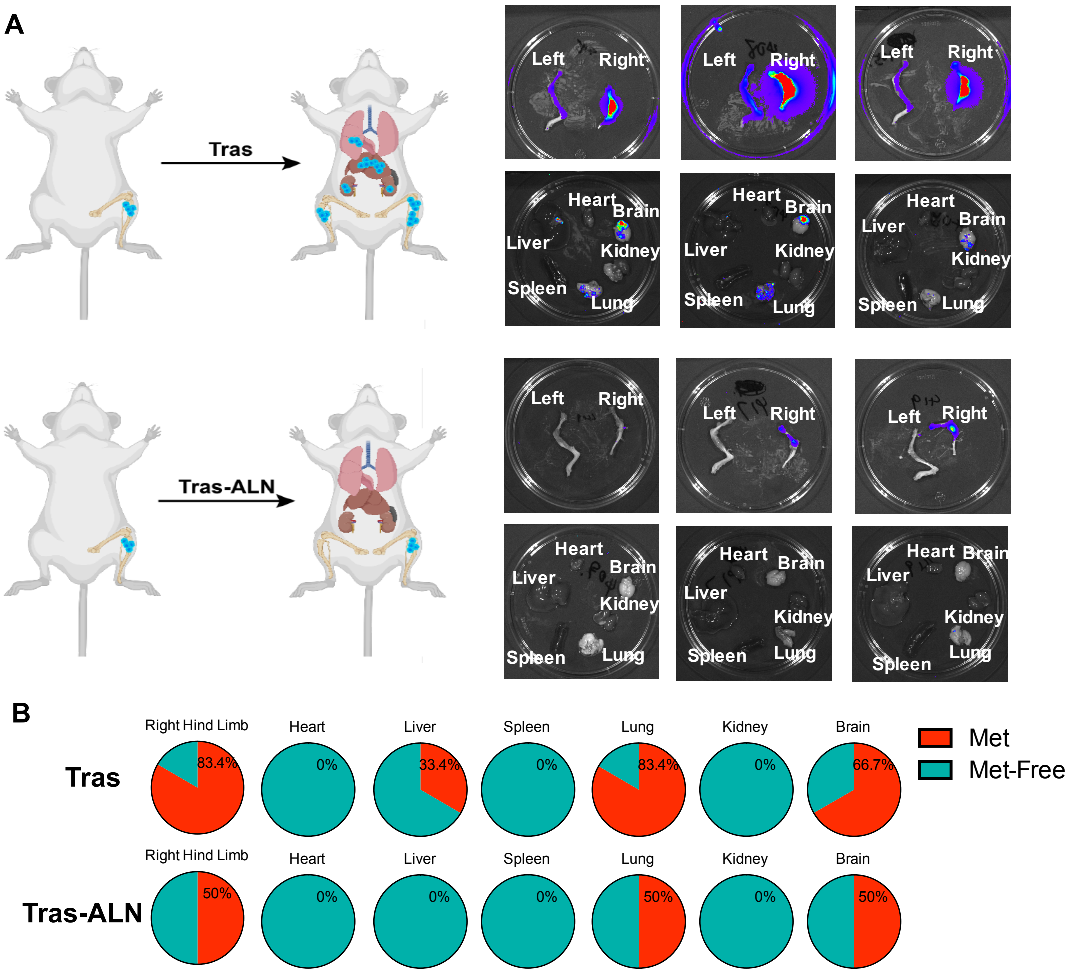

**Fig. S22 Tras-ALN effects on multi-organs metastases in MCF-7 cell lines.** (A) Metastases observed in various organs in mice treated with Tras or Tras-ALN. (B) Pie charts show the frequencies of metastasis observed in various organs in Tras and Tras-ALN treated groups.

**Table S1. Potency and cell-surface reactivity of Tras and Tras-ALN against breast cancer epithelial cell lines.**

| Cell Line | Her2 Expression | IC50 (µg/mL) | | MFI increase (fold) | |
|---|---|---|---|---|---|
| | | Trash | Tras-ALN | Tras | Tras-ALN |
| BT-474 | 3+ | 1.4 ± 0.9 | 2.3 ± 0.7 | 47.57 | 43.72 |
| SK-BR-3 | 3+ | ------- | ------- | 57.01 | 51.70 |
| MDA-MB-361 | 2+ | 57 ± 10 | 78 ± 21 | 23.50 | 31.30 |
| MDA-MB-468 | 0 | > 500 | > 500 | 1.01 | 1.06 |

Abbreviations: MFI, median fluorescence intensity. Binding was determined as the mean fold increase in median fluorescence over the PBS control.

**Table S2. Comparison of different treatment groups in multiple assays (MDA-MB-361 model).**

| Treatment | BLI | | Osteoclast activity | | | Body weight | |
|---|---|---|---|---|---|---|---|
| | Progression[a] | Fold-increase[b] | Osteoclast number[c] | Serum TRACP 5b[d] | Serum Calcium[e] | Progression[f] | Survival[g] |
| PBS vs ALN | NS | NS | * | NS | NS | NS | NS |
| PBS vs Tras | **** | **** | **** | NS | NS | NS | NS |
| PBS vs Tras-ALN | **** | **** | **** | *** | ** | NS | *** |
| ALN vs Tras | **** | **** | * | NS | NS | NS | NS |
| ALN vs Tras-ALN | **** | **** | **** | *** | * | NS | *** |
| Tras vs Tras-ALN | **** | **** | ** | * | ** | NS | * |

Abbreviations: ANOVA, analysis of variance; BLI, bioluminescence imaging; TRAP, tartrate-resistant acid phosphatase. [a]Signal intensity of BLI in whole body over the course of the experiment. [b]Signal intensity fold-increase of BLI after treatment (BLI of day 87/BLI of day 6). [C]Osteoclast number measurement from TRAP-stained tibia/femur sections at the end of experiment. [d]Serum TRACP 5b concentration at the end of experiment. [e]Serum calcium concentration at the end of experiment. [f]Body weight progression over the course of the experiment. [g]The mice BLI intensity over $10^7$ was considered to reach the endpoint. [a,b]were analyzed statistically by using a two-way repeated-measure ANOVA followed by Sidak's multiple comparisons test. [c,d,e,f] were analyzed by using a one-way ANOVA followed by Tukey's multiple comparisons test. [g]was analyzed by using a log-rank test. ****$P < 0.0001$, ***$P < 0.001$, **$P < 0.01$, *$P < 0.05$, NS represents $P > 0.05$.

**Table S3. Comparison of Tras and Tras-ALN groups in multiple assays (MCF 7 model).**

| Treatment | BLI | | Osteoclast activity | | Body weight | |
|---|---|---|---|---|---|---|
| | Progression[a] | Fold-increase[b] | Serum Calcium[c] | Serum TRACP 5b[d] | Increase[e] | Survival[f] |
| Tras vs Tras-ALN | **** | **** | * | * | NS | * |

Abbreviations: ANOVA, analysis of variance; BLI, bioluminescence imaging; TRAP, tartrate-resistant acid phosphatase. [a]Signal intensity of BLI in whole body over the course of the experiment. [b]Signal intensity fold-increase of BLI after treatment (BLI of day 68/BLI of day 6). [c]Serum calcium concentration at the end of experiment. [d]Serum TRACP 5b concentration at the end of experiment. [e]Body weight progression over the course of the experiment. [f]The mice BLI intensity over $10^7$ was considered to reach the endpoint. [a,b]were analyzed statistically by using a two-way repeated-measure ANOVA followed by Sidak's multiple comparisons test. [c,d,e]were analyzed by using a one-way ANOVA followed by Tukey's multiple comparisons test. [f]was analyzed by using a log-rank test. ****$P < 0.0001$, *$P < 0.05$, NS represents $P > 0.05$.
